# Multifaceted immune resistance landscapes in human oligodendrocytes protect against cytotoxic T cells and are dysregulated in MS brain cell subsets

**DOI:** 10.64898/2026.04.21.719872

**Authors:** Ayse Nur Menevse, Abir Hussein, Julian Sax, Antonio Sorrentino, Valentina Volpin, Nisit Khandelwal, Slava Stamova, Jasmin Mühlbauer, Nicole Schuch, Birgitta Ott-Rötzer, Antonia Engelhorn, Hannes Linder, Anchana Rathinasamy, Chih-Yeh Chen, Franziska C. Sanna, Alexander Wurzel, Leonard Bellersheim, Maria Xydia, Robert Lohmayer, Deniz Tümen, Karsten Gülow, Tillman Michels, Philipp Beckhove

## Abstract

Multiple sclerosis (MS) is a progressive neuroinflammatory demyelinating disease of the central nervous system (CNS) that remains incurable. Autoreactive myelin-specific T cells contribute to immunopathology by directly targeting and damaging oligodendrocytes *in situ*. In oligodendrocytes, several immune-modulatory functions have been described that can ameliorate immune damage. However, a systematic discovery of cell-intrinsic mechanisms that protect oligodendrocytes against T cell-derived cytotoxic mechanisms has not been performed. We used human MO3.13 oligodendrocytic cells and human antigen-specific cytotoxic T cells to conduct a high-throughput (HTP) RNAi-based screen with altogether 4155 genes to identify oligodendrocyte-intrinsic immune resistance genes (IRGs). The screen revealed 133 candidate IRGs. Among them, we validated 32, which exerted a strong immuno-protective phenotype. We studied IRG expression landscapes in human brain cell subsets from postmortem brain tissues of MS and control individuals. This revealed clustered expression of IRGs in a cell-type and oligodendrocyte subset-specific manner and differential IRG expression between MS patients and controls in distinct oligodendrocyte subclusters. ChEA3 analysis revealed cell type-specific expression of transcription factors that can drive expression of respective IRGs. Explorative molecular mode of action analyses of five selected IRGs, *STK11*, *KCNH8*, *ABCA2*, *SLC1A3* and *CHRNA1* revealed that these prevented death receptor-mediated apoptosis induced by T cell-derived cytotoxic molecules. In particular, they controlled TRAIL-induced apoptosis by suppressing JNK1 activation through interfering with several upstream pathways regulating metabolic, potassium, cholesterol, glutamate and acetylcholine homeostasis. In addition, STK11, ABCA2, and CHRNA1 regulated TRAIL-R2 surface expression contributing to increased TRAIL-sensitivity whereas KCNH8 expression in oligodendrocytes inhibited secretion of inflammatory cytokines by cytotoxic T cells. Taken together, we here demonstrate the existence of multiple co-expressed IRGs in human oligodendrocytes that regulate multifaceted mechanisms of T cell resistance and are dysregulated in oligodendrocyte subsets of MS patients.

## Introduction

Multiple sclerosis (MS) is a chronic demyelinating and neuroinflammatory autoimmune disorder of the central nervous system (CNS) and remains incurable.^1^ The major pathological hallmarks of MS are demyelinated lesions throughout the CNS that are highly infiltrated with T cells, B cells and myeloid cells which target oligodendrocytes and neurons.^2, 3^ Inflammatory factors secreted by infiltrated immune cells and CNS-resident microglia further exacerbate inflammation and demyelination.^4^ During MS progression, remyelination can cause transient clinical recovery following an inflammatory relapse.^5^ However, the remyelination rate depends on the maturation of oligodendrocyte progenitor cells (OPCs).^6^ Depending on the cell subset, immune cells can both promote OPC differentiation and myelination, as well as impair the myelin sheath by attacking mature oligodendrocytes.^7, 8^ Vice versa, oligodendrocytes can modulate immune cell activity by expressing pro- and anti-inflammatory factors such as, LRP1, ICAM-1, CXCL10, PD-L1, PD-L2 and TGF-β.^9–11^ Single-nucleus RNA sequencing (snRNA-seq) analysis of MS brain tissues identified distinct subtypes of oligodendrocytes with gene signatures associated with inflammation.^12, 13^ However, molecular mechanisms regulating the complex bi-directional interplay between oligodendrocytes and immune cells that are involved in the MS immunopathology are not well understood yet.

Here, we hypothesized that oligodendrocytes express a plethora of immune resistance genes (IRGs) that protect oligodendrocytes against autoreactive cytotoxic T cells. Dysregulation of IRGs in oligodendrocytes may lead to increased T cell activity and/or sensitize oligodendrocytes towards T-cell-mediated destruction, which may both contribute to demyelination and onset or severity of MS.

We established an *in vitro* human oligodendrocyte immune rejection model which is based on a human oligodendrocytic cell line MO3.13 and influenza antigen (Flu)-specific CD8^+^ T cells (FluTC) and conducted functional high-throughput (HTP) RNAi screening for a systematic identification of genes that confer immune resistance in oligodendrocytes. We show that oligodendrocytes express a broad panel of hitherto unknown IRGs. In the human brain, these are differentially expressed in subsets of oligodendrocytes and brain cell subsets and differentially (dys)regulated in oligodendrocyte subsets of MS patients. We further demonstrated that selected IRGs such as *STK11/LKB1*, *KCNH8*, *ABCA2*, *SLC1A3/EAAT1* and *CHRNA1* mediate MO3.13 oligodendrocyte-intrinsic resistance to direct antigen-specific T cell attack as well as to cytotoxic molecules secreted by activated T cells, by regulating a variety of TRAIL receptor-mediated apoptotic pathways and by inhibiting T cell activity. Thus, these genes might play a role in MS immunopathology and may provide starting points for future therapeutic avenues.

## Results

### Human MO3.13 oligodendrocytes express multiple IRGs that confer resistance to antigen-specific CD8^+^ T cell-mediated cytotoxicity

We established an *in vitro* co-culture model for T-cell mediated destruction of human oligodendrocytes that could be later adapted to the HTP siRNA-based screening platform previously developed by our group.^14–16^ Based on its high expression of oligodendrocyte markers such as myelin basic protein (MBP), we selected the immortalized human oligodendrocytic glial cell line MO3.13 as model target cell line.^17, 18^ We stably transfected MO3.13 with HLA-A2 and firefly-luciferase to enable recognition by HLA-A2 restricted cytotoxic T cells and to quantify MO3.13 oligodendrocyte killing (Figure 1A-B). Gene silencing by siRNA transfection was consistently efficient in MO3.13-A2-Luc cells as evidenced by complete cell death 72 h post transfection with a siRNA cocktail targeting essential cell survival genes (“cell death” siRNA, siCD) (Figure 1C) and by significant downregulation of *PDL1* mRNA and surface protein expression following siRNA targeting of PD-L1 in MO3.13-A2-Luc (Figure 1D–E).

**Figure 1:**
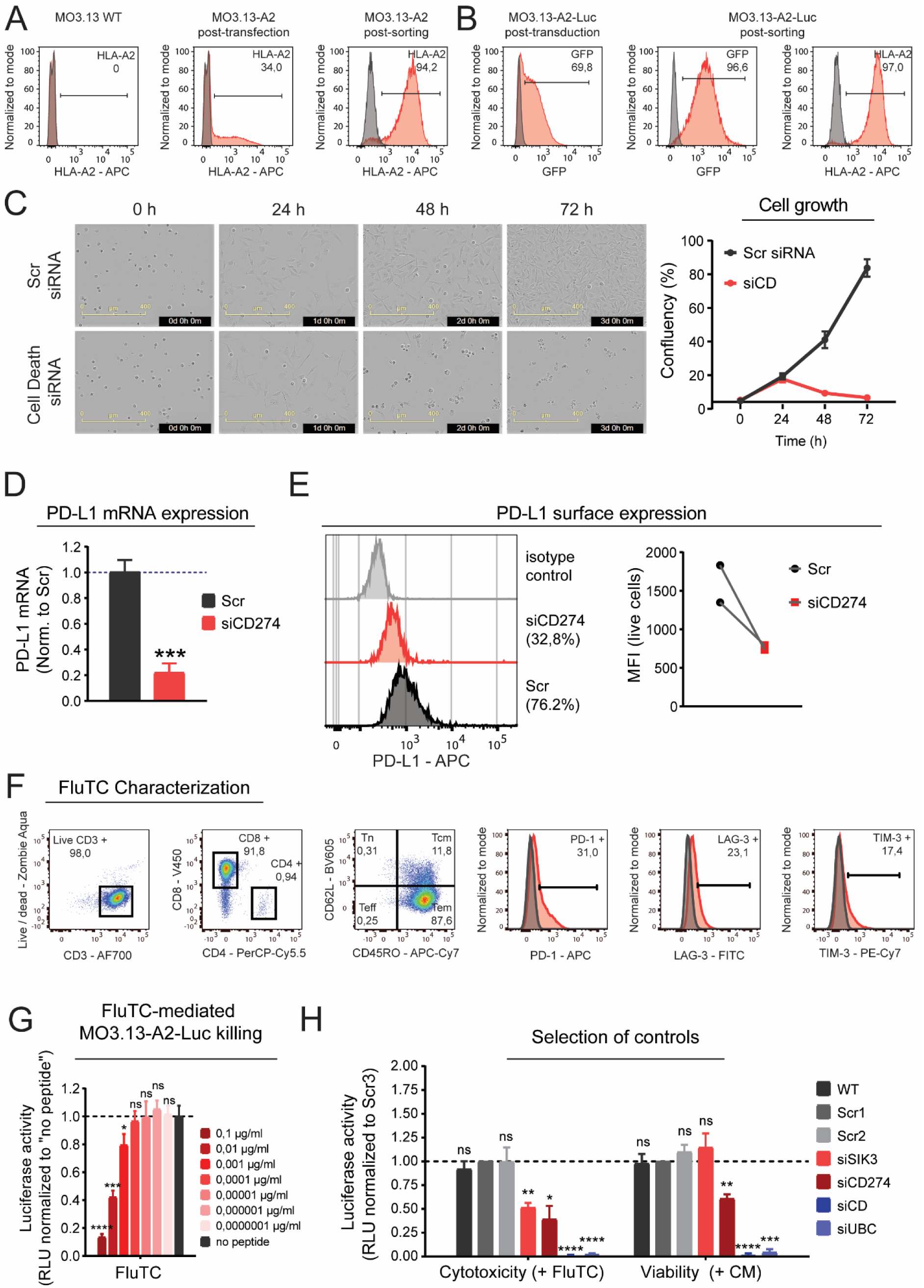
Establishment of an *in vitro* co-culture model for MS for the HTP screen. **(A-B)** Generation of HLA-A2^+^ Luciferase^+^ human oligodendrocytic cell line MO3.13-A2-Luc. (A-B) FACS analysis of the (A) HLA-A2 expression on wildtype (WT) MO3.13 cells (left), MO3.13-A2 cells 2 weeks after transfection with HLA-A2 expression vector (middle), and MO3.13-A2 cells after the FACS sorting for HLA-A2 (right). (B) GFP expression on MO3.13-A2-Luc cells after lentiviral transduction (left), GFP expression on FACS sorted MO3.13-A2-Luc cells (middle) and HLA-A2 expression on MO3.13-A2-Luc cells (right). Grey histograms represent the isotype control and untransduced cells for the analysis of HLA-A2 and GFP expression respectively. **(C-E)** Optimization of siRNA reverse transfection protocol for MO3.13-A2-Luc cells. (C) Real-time live cell imaging of MO3.13-A2-Luc cells transfected either with a non-targeting siRNA control (Scr) or with a siRNA cocktail targeting genes essential for cell survival (cell death siRNA, siCD). Transfected cells were imaged via the IncuCyte SX5 system for 72 h and real-time % cell confluency was quantified. (D) RT-qPCR analysis of *PDL1* mRNA expression in MO3.13-A2-Luc cells transfected either with Scr siRNA or pool of 4 non-overlapping siRNAs targeting *PDL1* (siCD274). Results are presented as fold change compared to the Scr after *β-actin* mRNA normalization. (E) FACS analysis of PD-L1 surface expression on MO3.13-A2-Luc cells transfected either with Scr or PD-L1 specific siRNA pool. Left: representative histograms indicating % PD-L1 expression, right: mean fluorescent intensity (MFI) of PD-L1 from 2 independent FACS data. **(F)** Phenotypic characterization of FluTC. The expression of CD4 and CD8 was determined on CD3^+^ FluTC, whereas the expression of effector and memory markers CD45RO and CD62L and co-inhibitory immune checkpoint molecules PD-1, LAG-3 and TIM-3 was determined on CD3^+^ CD8^+^ FluTC. (Tn: naïve T cells, Tcm: central memory T cells, Teff: terminal effector T cells, Tem: effector memory T cells). Grey histograms represent the isotype control. **(G)** Luciferase-based cytotoxicity assay to optimize the co-culture conditions for MO3.13-A2-Luc and FluTC. MO3.13-A2-Luc were pulsed with serial dilutions of the flu-peptide and co-cultured with FluTC. After 20 h of co-culture, remaining relative luciferase units (RLU) were measured. RLU measured in pulsed samples were normalized to RLU of unpulsed (no peptide) control. Statistical significance was calculated compared with no peptide control. **(H)** Selection of cytotoxicity and viability controls for the HTP screen. MO3.13-A2-Luc cells were transfected with the indicated siRNAs. After 72 h of transfection cells were pulsed with 0,01 µg/ml flu-peptide and co-cultured either with FluTC or treated with control T cell medium (CM). 20 h following co-culture, remaining RLU was measured. The RLU of each sample was normalized to the Scr1 control in the cytotoxicity (FluTC) or viability (CM) setting to determine the impact of gene knockdown. (A-G) Representative data of at least 2 independent experiments. (H) Cumulative data of three independent experiments each performed with four replicates per sample. (D, G-H) Graphs show mean +/- SD. p-values were calculated using two-tailed student’s t-test. * = p < 0.05, ** = p < 0.01, *** = p < 0.005, **** = p < 0.001.

In MS patients, a wide range of myelin-derived autoantigenic peptides have been identified whose immunogenicity varies depending on the peptide sequence and the patient’s HLA haplotype.^19^ To eliminate this variability, we loaded MO3.13-A2 cells with HLA-A2-restricted peptide derived from the Influenza matrix protein M1 (flu-peptide) as a model autoantigen.^20^ To mimic myelin-reactive cells in MS patients and study antigen-specific CD8^+^ T cell-mediated killing of oligodendrocytes, Influenza matrix protein M1 (flu-peptide)-specific T cells (FluTC) were generated from PBMCs of HLA-A2+ healthy donors by repetitive antigen stimulation as described before (Supp. Fig. 1).^20^ FluTC constitute more than 90% of the generated T cell cultures and display effector memory (Tem) and central memory (Tcm) phenotype (Supp. Fig. 1A, Figure 1F). FluTC expressed low levels of activation/exhaustion markers PD-1, LAG-3, and TIM-3 and secreted high levels of IFNγ and granzyme B upon co-culture with peptide loaded MO3.13-A2-Luc cells indicating their strong cytotoxic potential and capacity for further activation (Figure 1F, Supp. Fig. 1B).^21^ FluTC-mediated antigen-specific kill of MO3.13-A2-Luc cells was analyzed by pulsing MO3.13-A2-Luc with diluting concentrations of flu-peptide (Figure 1G). We chose 0,01 µg/ml as optimal peptide concentration to induce an intermediate FluTC-mediated cytotoxicity (Figure 1G).

The screen consisted of cytotoxicity and viability settings where siRNA-transfected and flu-peptide-pulsed MO3.13-A2-Luc cells were co-cultured either with FluTC (*cytotoxicity setting*) or treated with control T cell medium (CM) (*viability setting*). Following 20 h of co-culture, luciferase activity was measured, which is proportional to the amount of remaining living cells in each well. The *cytotoxicity setting* allows the identification of genes whose knockdown increases or decreases T cell-mediated MO3.13 oligodendrocyte killing, whereas the *viability setting* allows the exclusion of genes whose knockdown has an impact on cell viability *per se*. Two different non-targeting siRNA sequences (Scr1 and Scr2) that displayed similar phenotypes as the untransfected wild type control (WT) were included as negative controls (Figure 1H). To identify positive controls for the cytotoxicity setting, several established IRGs were tested among which the knockdown of SIK3 in MO3.13-A2-Luc cells resulted in increased FluTC-mediated cytotoxicity without an impact on cell viability *per se* (Figure 1H). Downregulation of *PDL1*/*CD274* also improved FluTC cytotoxicity while imposing a slight impact on MO3.13-A2-Luc viability (Figure 1H). As positive controls for the viability setting, a cell death siRNA cocktail and ubiquitin C (UBC) siRNA were included as these efficiently induced cell death (Figure 1H).

For the screen we used a siRNA library targeting 4155 genes which comprised the entire surfaceome (3702 membrane-bound molecules (89%)) and 453 kinases and metabolic proteins (11%) (Supp. table 1) and which covered 50 of the most established immune checkpoint genes (Supp. table 3). Each gene was targeted by a pool of 4 non-overlapping siRNAs (Supp. table 1). Transfected MO3.13-A2-Luc cells were either co-cultured with FluTC (cytotoxicity setting) or cultured in plain T cell media (viability setting) and luciferase readout was performed 21 h post co-culture. Plate normalization was performed as a first step of the data analysis, to exclude inter-plate variability during the readout (Figure 2A). As an indicator for the robust technical performance of the screen, the Pearson correlation coefficient (*r*^2^) among the replicates was calculated as *r*^2^ = 0.94 and *r*^2^ = 0.96 for the cytotoxicity and viability settings respectively (Figure 2B). Reliable performance of the non-targeting siRNA controls (WT, Scr1, Scr2), positive immune resistance controls (siSIK3, siCD274), apoptosis resistance control (siCASP3) and viability control (siCD) are demonstrated in Figure 2B. To define IRGs, we transformed the RLU values of each gene knockdown into z-scores, which are defined by the number of standard deviations between the corresponding data point and the mean RLU of the plate (Figure 2C) multiplied by −1 (for simplification of analysis). Thereby, positive z scores indicate reduced luciferase intensity. Viability and cytotoxicity scores of each gene were plotted on quadrant plots. Candidate IRGs (HITs) with high cytotoxicity and low viability scores are depicted in Figure 2C (rectangle). To rank candidate IRGs according to their immuno-protective impact, we calculated differential scores; the difference between the cytotoxicity and the viability z-scores using the local regression (LOESS) rank (Figure 2D). The primary screen revealed 133 IRGs that were ranked higher than the strongest performing positive control SIK3 (Supp. table 2). We evaluated the performance of 50 well-established immune checkpoint genes that were covered in the library (Supp. table 3). Several of them were among the 133 top-ranked IRGs, such as *CEACAM6*^22^ (Carcinoembryonic antigen-related cell adhesion molecule 6), *NT5E*^23^ (5’-nucleotidase, CD73), *HLA-G*^24^ (HLA class I histocompatibility antigen, alpha chain G), *SIGLEC1*^25^ (Sialic acid-binding Ig-like lectin 1), and *OLR1*^26^ (Oxidized Low-Density Lipoprotein Receptor 1) (Figure 2C & D).

**Figure 2:**
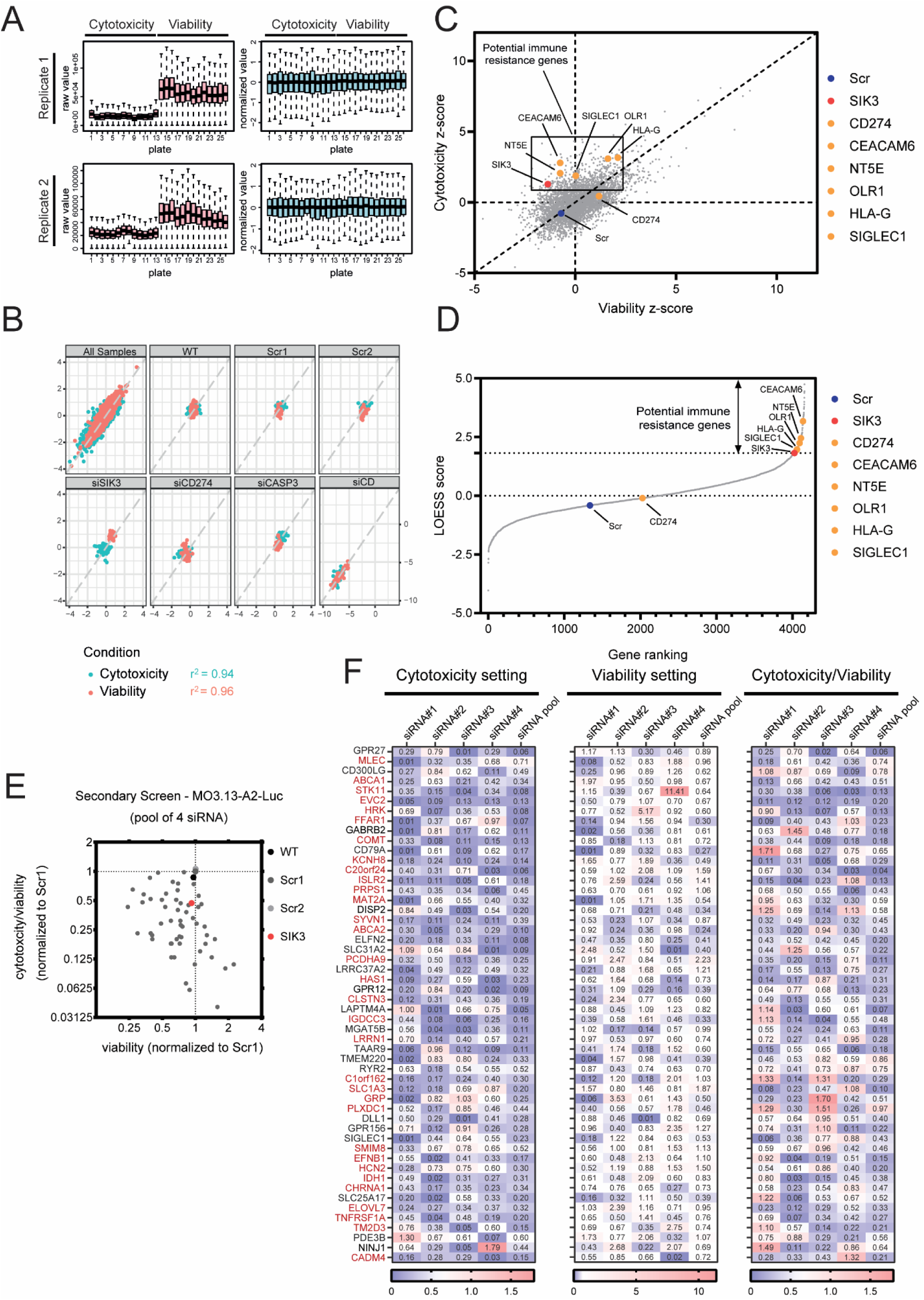
HTP screen to identify novel immune resistance genes in MO3.13 oligodendrocytes. **(A-B)** Performance of the controls. (A) Left panel: Raw luciferase activity (RLU) was measured for each well of 52 x 384-well plates. Upper and lower panels show replicate 1 & 2 respectively. Right panel: To exclude inter-plate variability, RLUs were normalized using the following formula: Normalized RLU =**log_2_ *x/M***, where ***x*** is the raw RLU from each well and ***M*** is the median RLU value in each plate. For each replicate set, plates from 1 to 13 were co-cultured with FluTC at E:T = 20:1 (Cytotoxicity), while plates from 14 to 26 were cultured with plain T cell media (Viability) (B) Performance of the controls in the HTP-screen. Dot plot shows normalized RLUs after transfection of MO3.13-A2-Luc cells with several control siRNAs. Technical replicates were plotted against each other (x-axis: replicate 1; y-axis: replicate 2). Blue dots: cytotoxicity setting (with FluTC). Red dots: viability setting (plain medium). Pearson correlation (r^2^) amongst the 2 replicate values was calculated for each setting (cytotoxicity setting: r^2^ = 0,94; viability setting: r^2^ = 0,96). **(C-D)** Performance of the HTP screen for the identification of novel MS-associated IRGs. (C) Overview of the results from the primary screen where the quadrant plot depicts the z-scores of the median-normalized luciferase intensity of transfected MO3.13-A2-Luc cells after co-culture with FluTC (cytotoxicity score) or with culture medium (viability score), using a siRNA library of 4155 genes plus control genes. The cytotoxicity score indicates the influence of the gene knockdown on FluTC-mediated killing. Positive values indicate decreased tumor cell viability. Potential IRGs with high cytotoxicity and low viability scores are depicted in a box. (D) Gene ranking diagram showing the differential score between the cytotoxicity and viability z-scores using the local regression (LOESS) rank. Potential IRGs are represented in the ascending arm of the rank. Scores of Scr control and known IRGs *PDL1*/*CD274*, *CEACAM6*, *NT5E*, *OLR1*, *HLA-G*, *SIGLEC1* and *SIK3* are highlighted in the graph. **(E-F)** Results of the secondary screen with MO3.13-A2-Luc cells. A deconvoluted siRNA library consisting of pool of 4 non-overlapping siRNAs as well as single siRNAs targeting selected 52 HITs from the primary screen was used to transfect MO3.13-A2-Luc cells. 72 h post transfection, cells were co-cultured either with FluTC or cultured in T cell media as performed in the primary screen. RLU values from cytotoxicity and viability settings were measured after 24 h of co-culture and normalized to the Scr control. (E) Dots depict the average normalized RLU values obtained from cells transfected with the pool of 4 non-overlapping siRNAs. (F) Heatmap indicating the average normalized RLU values of MO3.13-A2-Luc cells transfected either with single siRNA or pool of 4 siRNAs measured in cytotoxicity and viability settings. HITs with a stronger immune resistance profile than SIK3 were marked in red. (E-F) Each siRNA transfection was performed in quadruplicates.

Among the 133 HITs we selected 52 which i) were expressed within the human brain, ii) whose role in immune regulation has not been well-established and iii) excluded key components of essential cellular maintenance machineries or olfactory/taste receptors (GPCR) (Supp. table 4). To validate the immune resistance phenotype of these HITs, we performed a secondary screen to exclude off-target effects. To this end we deconvoluted the siRNA pools into four single non-overlapping siRNAs per gene (Figure 2E & F). The cytotoxicity and viability RLU values of each siRNA condition were normalized to the RLU values of Scr in each setting. Downregulation of SIK3 as the positive control IRG revealed values for cytotoxicity of 0,43, viability of 0,92 and a cytotoxicity/viability (c/v) ratio of 0,47. Accordingly, we considered IRGs with cytotoxicity ratio < 0,43 and/or c/v ratio < 0,47 and viability ratio > 0,6 for at least two single siRNA sequences as “IRGs with stronger immune resistance phenotype than SIK3”. 32 HITs out of 52 matching these criteria showed again a strong (>SIK3) immune resistance function and were selected for further analysis (Figure 2F, Supp. table 5). The majority of them (21) are expressed on the outer cell membrane, 6 on the intracellular membranes of the ER, mitochondria or lysosomes, 4 are cytoplasmic enzymes (STK11, PRPS1, MAT2A, IDH1), and 1 IRG is a secreted 27aa peptide (GRP). Among 27 IRGs localized to the cell/organelle membrane, 15 are single-pass and 12 are multi-pass membrane proteins.^27^

### Human oligodendrocytes express a distinct cluster of co-regulated IRGs with subpopulation and MS-associated signature

To examine the expression of the identified 133 IRGs in primary human oligodendrocyte subsets *in situ* and to identify any potential MS-associated differential IRG expression, we analyzed a public single-nucleus RNA-Seq (snRNA) dataset comprising 17.799 nuclei derived from post-mortem brain tissues of four patients with progressive MS and five control individuals (Figure 3, Supp. Fig. 2, Supp. table 6).^12^ Average gene expression was assessed in the following brain-resident cell clusters as recently defined and annotated by Jäkel *et*. *al*^12^: oligodendrocyte progenitor cells (OPCs), committed oligodendrocyte precursors (COPs), inflammation associated immune oligodendroglia (ImOLG), six mature oligodendrocyte subclusters (Oligo1-6), two astrocyte subclusters, microglia-macrophages, macrophages, immune cells, five neuron subclusters (Neuron1-5), two endothelial cell subclusters, pericytes and vascular smooth muscle (VSM) cells (Figure 3A, Supp. Fig. 2A). Apart from *SIK3*, 32 HITs out of 133 were detected among the differentially expressed marker genes of at least one cell cluster and are displayed in Figure 3A and in Supp. Fig. 2A. Our analysis revealed that each defined major cell type (OPCs, COPs, ImOLGs, oligodendrocytes, astrocytes, microglia, macrophages, neurons, endothelial cells, pericytes, VSM cells) differentially upregulated clearly distinct panels of multiple IRGs. ImOLGs displayed an IRG profile between COPs and mature oligodendrocytes, while sharing IRGs with astrocytes and microglia/macrophage (microglia/mac) clusters (Figure 3A, Supp. Fig. 2A). Those IRGs that were differentially expressed in individual oligodendrocyte subclusters compared to all other cell clusters are depicted in Figure 3B and in Supp. Fig. 2B. Expression levels of validated strong IRGs (Figure 2F) and *SIK3* are also highlighted in volcano plots (Figure 3B-C, Supp. Fig. 2B-C). *ABCA2*, *KCNH8* and *SIK3* were consistently upregulated in ImOLGs and most oligodendrocyte clusters (Figure 3B, Supp. Fig. 2B). On the other hand, *SLC1A3* was significantly upregulated only in ImOLGs and COPs but downregulated in most mature oligodendrocyte clusters (Figure 3B, Supp. Fig. 2B). As oligodendrocyte heterogeneity and composition can be altered in patients with MS^12^, we compared the expression of IRGs in different oligodendrocyte subclusters in MS patients and controls (Figure 3C, Supp. Fig. 2C). Among the 32 validated IRGs, *ABCA2* expression was consistently increased across all oligodendrocyte subclusters in MS patients, with significant upregulation detected in the majority of them (Oligo1/4/5/6) (Figure 3C, Supp. Fig. 2C). In contrast, *KCNH8* and *COMT* were significantly downregulated in Oligo2 and Oligo5 subclusters of MS patients respectively (Figure 3C). Within the mature oligodendrocyte subclusters *KCNH8*, *ABCA2* and *SIK3* were expressed by the majority of cells, whereas *SLC1A3* was expressed by the majority of progenitor and ImOLG cells (Figure 3D).

**Figure 3:**
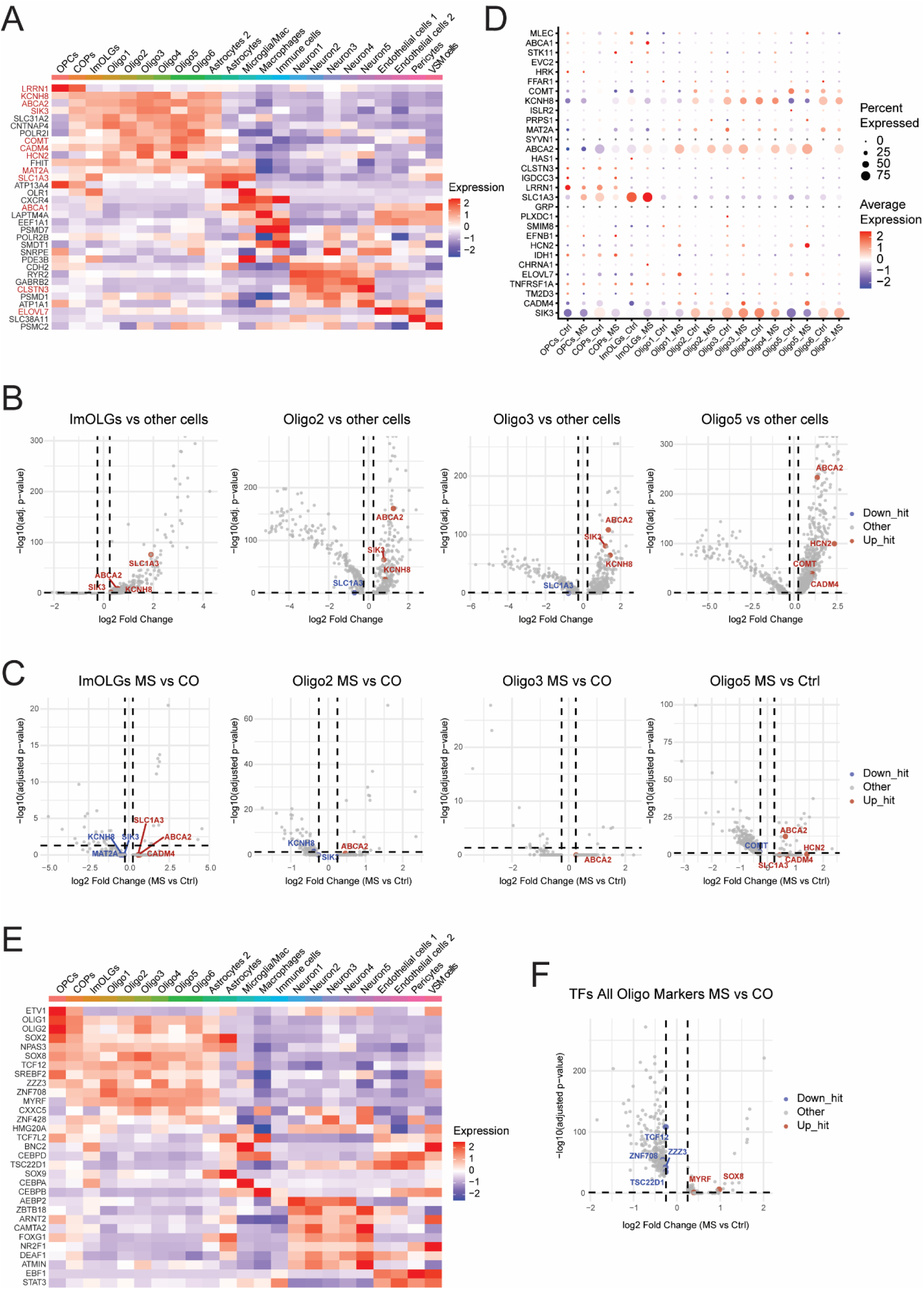
Human oligodendrocytes co-express distinct sets of IRGs with a subpopulation and MS-associated signature. Gene expression of HITs identified in the primary screen was assessed among all identified populations of cells from snRNA-seq data from Jäkel *et al.* as annotated by the authors.^12^ **(A)** Only HITs that are among differentially expressed marker genes of different cell clusters are shown in an average expression heatmap. Validated IRGs with strong immuno-protective phenotype are marked in red. **(B-C)** Volcano plots depicting (B) DEGs of each individual oligodendrocyte subcluster versus all other cell types in MS and control samples combined and (C) DEGs of each individual oligodendrocyte subcluster from MS versus control samples. Only the IRGs with a strong immune resistance phenotype and SIK3 were labeled. **(D)** The average gene expression of IRGs identified and validated in the secondary screen between MS and control oligodendrocyte subpopulations are visualized in a dot plot depicting scaled average expression levels and the percentage of cells expressing the respective gene across every population (dot size). **(E)** Top 100 TFs that are among the differentially expressed marker genes of different cell clusters are shown in an average expression heatmap. **(F)** Volcano plots depicting differentially expressed TFs between MS and control samples. (B, C, F) Upregulated and downregulated IRGs/TFs were marked in red and blue respectively. Horizontal and vertical dashed lines indicate log_2_ fold-change thresholds (±0,25) and adjusted p-value cutoff −log_10_(0,05). Both heatmaps and dot plot display expression levels indicated by a color gradient ranging from blue to white to red.

We then explored the transcription factors (TFs) that are predicted to drive the expression of the validated IRGs within the various oligodendrocyte cell clusters. We submitted 32 IRGs alongside SIK3 to the ChIP-X Enrichment Analysis Version 3 (ChEA3) web-server and obtained the average integrated rank (mean rank) list of the transcription factors upstream of multiple overlapping target IRGs (Supp. table 7).^28^ Remarkably, among the top 10 TFs we found MYRF^29^, OLIG2^30^, NKX6-2^31^, SOX8^32^, SOX2^33^ which are known crucial regulators of oligodendrocyte-specific gene expression. Other oligodendrocyte-associated TFs such as RXRA^34^, TCF7L2^35^, SREBF1^36^, OLIG1^37^, SREBF2^38^, SOX10^39^ were ranked among the top 50 TFs (Supp. table 7). To determine a broader range of oligodendrocyte-enriched TFs, we selected the 100 top ranked TFs identified by ChEA3 and analyzed their average expression in different brain cell populations from the snRNA-seq dataset (Figure 3E, Supp. Fig. 2D). This revealed additional TFs that were significantly enriched among oligodendrocyte subclusters with yet limited or uncharacterized functions in oligodendrocytes such as NPAS3, TCF12^40^, ZZZ3, ZNF708^41^ (Figure 3E, Supp. Fig. 2D).

In line with the co-expression profile observed in oligodendrocyte subclusters (Figure 3A-D, Supp. Fig. 2A-B), *ABCA2* and *KCNH8* were predicted to be co-regulated by MYRF, NKX6-2, SOX8 and ZNF708 (Supp. Table 7). Consistent with the marker and IRG expression profile, ImOLGs displayed a TF expression profile similar to both oligodendrocytes as well as to astrocytes-microglia/macrophage populations (Figure 3E, Supp. Fig. 2D). Notably, the ImOLG-enriched IRG SLC1A3 (Figure 3B) was predicted to be regulated by both oligodendrocyte-enriched TFs such as OLIG1, OLIG2, SOX2, NPAS3, SOX8, TCF12 and SREBF2, as well as by astrocyte-microglia/macrophage enriched TFs such as BNC2, CEBPD and SOX9 (Supp. table 7, Figure 3E, Supp. Fig. 2D). Although we could not detect a prominent *STK11* and *CHRNA1* expression in the snRNA-seq dataset, potentially due to either technical limitations or low nuclear retention (Supp. Fig. 2E),^42^ they were predicted to be co-regulated by OLIG2, RXRA and TCF12, and other oligodendrocyte-associated TFs which also drive *ABCA2, KCNH8* and/or *SLC1A3* expression (Supp. table 7). We subsequently assessed the differential expression of transcription factors predicted to regulate IRG expression in MS patients (Figure 3F). The expression of oligodendrocyte-associated TFs MYRF and SOX8 were significantly upregulated in MS samples (Figure 3F), which can be the underlying reason for upregulated *ABCA2* expression in oligodendrocytes of MS patients (Figure 3C, Supp. fig. 2C).

Based on the performance in the primary and secondary screen, snRNA-seq expression data and ChEA3 analysis, we selected five novel IRGs, namely *STK11* (Serine/threonine-protein kinase STK11), *KCNH8* (Voltage-gated delayed rectifier potassium channel KCNH8), *ABCA2* (ATP-binding cassette sub-family A member 2), *SLC1A3* (Excitatory amino acid transporter 1) and *CHRNA1* (Acetylcholine receptor subunit alpha) for further functional characterization to determine molecular mechanisms through which they confer immune resistance in oligodendrocytes.

### Identified IRGs protect MO3.13 oligodendrocytes against TRAIL-induced apoptosis by controlling JNK activation

We conducted exploratory MoA analyses in both undifferentiated and differentiated MO3.13-A2-Luc cells which showed stable expression of *STK11*, *KCNH8*, *ABCA2*, *SLC1A3* and *CHRNA1* whereby only the expression of *KCNH8* was moderately decreased upon maturation (Figure 4A). Transfection of MO3.13-A2-Luc cells with IRG-targeting siRNA pool resulted in significant gene knockdown (Figure 4B) and in accordance with the screen results, IRG-deficient MO3.13 oligodendrocytes underwent stronger FluTC-mediated lysis compared to MO3.13 oligodendrocytes transfected with non-targeting siRNA pool (Supp. Fig. 3A).

**Figure 4:**
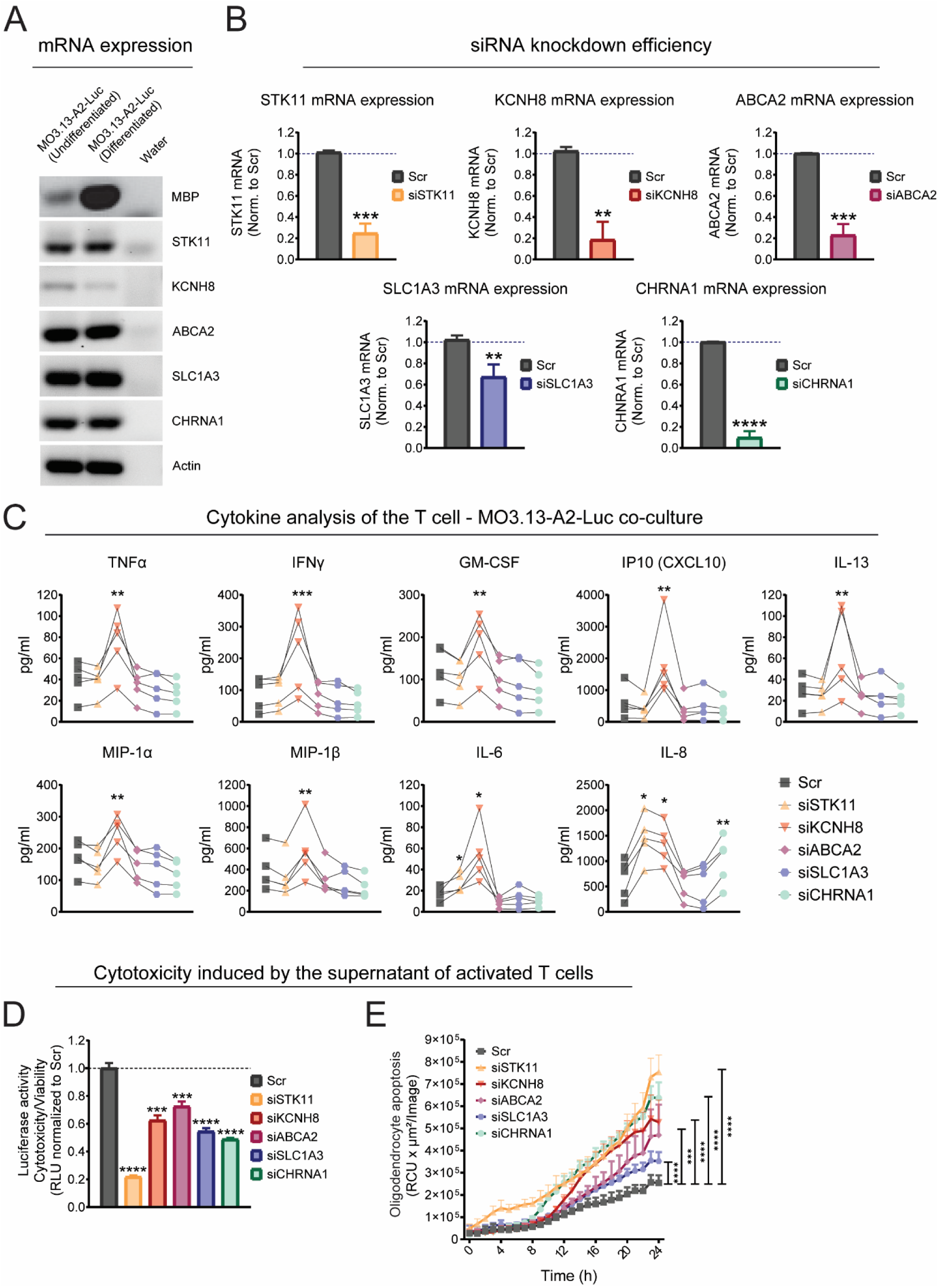
The impact of selected IRGs on T cell function and MO3.13 oligodendrocyte resistance against cytotoxic molecules secreted by activated T cells. **(A)** Expression of selected HITs in MO3.13-A2-Luc cells. Conventional PCR was performed to detect gene expression of selected HITs in undifferentiated and differentiated MO3.13-A2-Luc cells. Myelin basic protein (MBP) expression was used as positive control for MO3.13 oligodendrocyte differentiation and β-actin was used as house-keeping gene. **(B)** RT-qPCR analysis of *STK11, KCNH8, ABCA2, SLC1A3* and *CHRNA1* mRNA expression in MO3.13-A2-Luc cells transfected either with Scr siRNA or pool of 30 siRNAs targeting corresponding genes. Results are presented as fold change compared to the Scr after β-actin mRNA normalization. **(C)** Luminex-based cytokine analysis of the FluTC – MO3.13-A2-Luc cell co-cultures. MO3.13-A2-Luc cells transfected either with Scr siRNA or IRG-specific pool of 30 siRNAs and co-cultured with FluTC for 24 h. TNFα, IFNγ, GM-CSF, CXCL10, IL-13, MIP-1α, MIP-1β, IL-6 and IL-8 levels were depicted. Each line represents an independent experiment; values indicate the average of triplicates. **(D-E)** Selected IRGs protect MO3.13 oligodendrocytes against cytotoxic molecules secreted by activated T cells. MO3.13-A2-Luc cells transfected either with Scr siRNA or IRG-specific siRNA pool and treated with the supernatant of CD3/CD28-activated FluTC. (D) MO3.13 oligodendrocyte survival was determined by luciferase-based cytotoxicity assay. (E) Real-time cytotoxicity assay (IncuCyte® SX5 System) to analyze activated supernatant-induced MO3.13 oligodendrocyte apoptosis over 24 h in MO3.13-A2 cells. Cells were transfected and treated as in (D). Incucyte® Cytotox-Red Dye was added as an indicator of apoptosis. The graph shows total red object integrated intensity per well (RCU x µm²/Image). (A, D-E) Representative data of at least three independent experiments. (B-C) Cumulative data of three and five independent experiments respectively. Values represent (B, D-E) the mean ± SD, (C) mean. P-value was calculated using paired two-tailed Student’s t-test (* = p < 0.05, ** = p < 0.01, *** = p < 0.005, **** = p < 0.001).

Oligodendrocytes can exploit several protective mechanisms to evade autoreactive-immune cell attack.^43, 44^ These encompass the suppression of immune cell activity by expressing immune checkpoint molecules such as PD-L1 and PD-L2, and improvement of cell-intrinsic resistance against cytotoxic factors such as NF-*κ*B activation and cFLIP upregulation.^43, 44^ We studied the production of a broad array of cytokines by FluTCs upon their co-culture with MO3.13-A2-Luc cells. Silencing of *KCNH8* in MO3.13 oligodendrocytes significantly increased the concentration of TNFα, IFNγ, GM-CSF, CXCL10, IL-13, MIP-1α, MIP-1β, IL-6 and IL-8 (but not MCP-1) in the co-culture supernatant, while silencing of the other four IRGs did not affect cytokine secretion by FluTCs (Figure 4C, Supp. Fig. 3B). However, downregulation of *STK11* and *CHRNA1* elevated IL-8 expression in MO3.13-A2-Luc cells alone (Supp. Fig. 3C).

As *STK11*, *ABCA2*, *SLC1A3* and *CHRNA1* did not regulate FluTC activity, we investigated whether these IRGs can protect MO3.13 oligodendrocytes against soluble cytotoxic molecules secreted by TCs upon activation. Knockdown of all five IRGs increased MO3.13 oligodendrocyte lysis and apoptosis upon treatment with the supernatant of activated FluTC (Figure 4D, E). Thus, STK11, KCNH8, ABCA2, SLC1A3 and CHRNA1 mediate MO3.13 oligodendrocyte-intrinsic resistance towards cytotoxic molecules released by activated TCs, whereby KCNH8 has an additional role in controlling T cell activation as well.

We further investigated the distinct T-cell-derived cytotoxic molecules against which the individual IRGs mediate MO3.13 oligodendrocyte resistance. As the supernatant of activated T cells, including FluTC, contains TNFα, IFNγ, FASL, and TRAIL (Supp. Fig. 3D)^20^, we analyzed the expression of their cognate pro-apoptotic receptors on MO3.13-A2-Luc cells (Figure 5A). Our analysis revealed a low expression of IFNG-R1, TNF-R1 and FAS and a strong expression of TRAIL-R2, whereas TNF-R2 and TRAIL-R1 were not expressed (Figure 5A). Due to involvement of TRAIL and TRAIL-R in MS susceptibility and immunopathology^45^, we hypothesized that the selected IRGs may protect MO3.13 oligodendrocytes towards TRAIL-mediated apoptosis. Silencing of *STK11*, *ABCA2* and *CHRNA1* increased TRAIL-R2 surface expression in MO3.13-A2-Luc cells, indicating an increased sensitivity against TRAIL-mediated cytotoxicity (Figure 5B). Although the expression of KCNH8 and SLC1A3 did not change TRAIL-R2 levels, silencing of all five IRGs as well as *SIK3* increased apoptosis in MO3.13-A2-Luc cells upon TRAIL treatment compared to IRG proficient cells (Figure 5C, Supp. Fig. 4).

**Figure 5:**
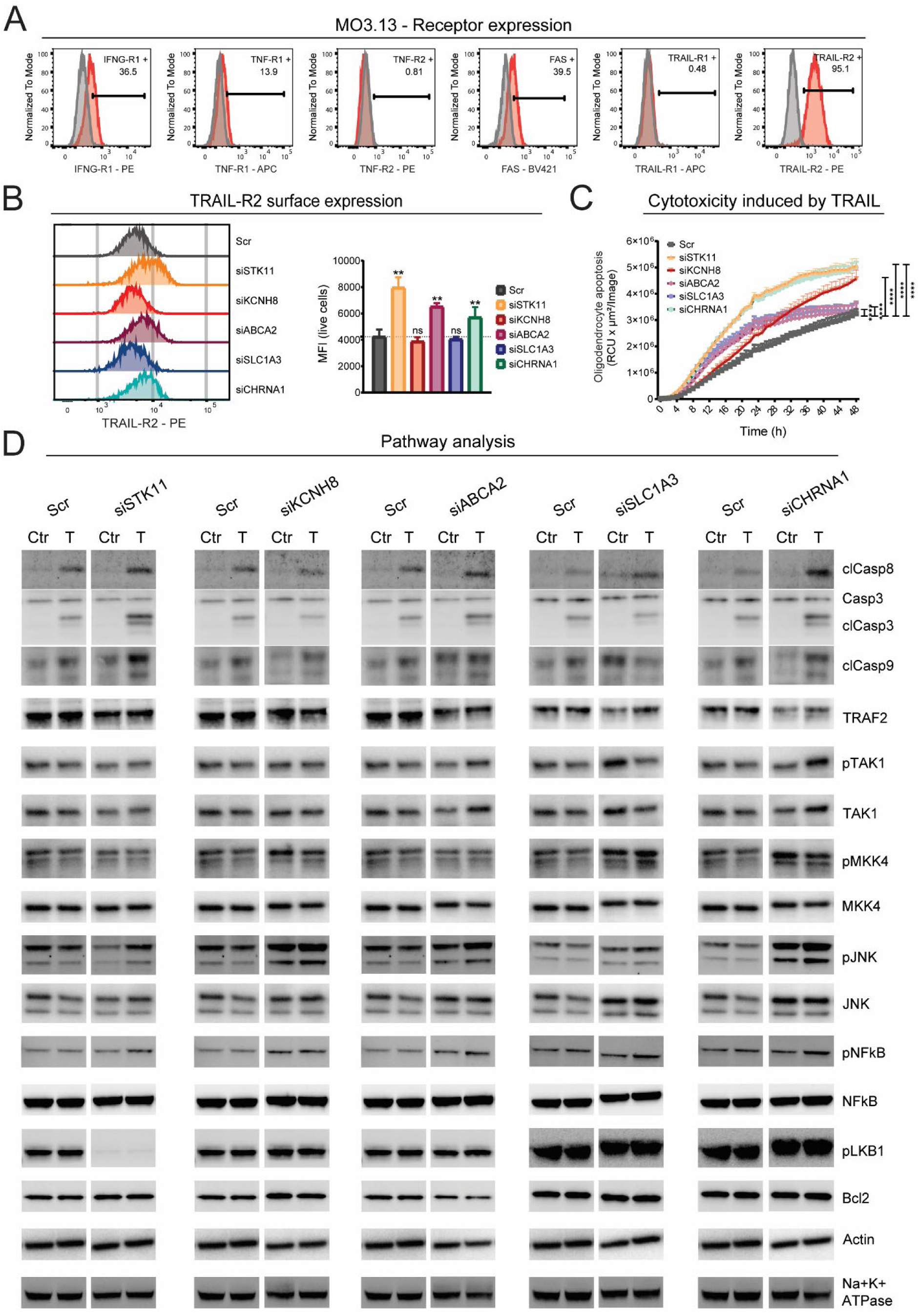
Selected IRGs protect MO3.13 oligodendrocytes from TRAIL-induced apoptosis. **(A)** FACS analysis to determine the surface expression of IFNG-R1, TNF-R1, TNF-R2, FAS, TRAIL-R1, and TRAIL-R2 in MO3.13-A2-Luc cells. Grey histograms represent the isotype control. **(B)** Impact of IRG knockdown on TRAIL-R2 surface expression in MO3.13-A2-Luc cells. Left panel: Representative overlay of histograms to compare TRAIL-R2 expression on IRG +/- MO3.13-A2-Luc cells. Right panel: Compiled data of mean fluorescence intensity (MFI) for TRAIL-R2 expression. **(C)** Real-time cytotoxicity assay (IncuCyte® SX5 System) to analyze TRAIL-induced apoptosis over 48 hours in IRG +/- MO3.13-A2-Luc cells. Incucyte® Cytotox-Red Dye was added to the transfected MO3.13 oligodendrocytes together with TRAIL treatment as an indicator of apoptosis. The graph shows total red object integrated intensity per well (RCU x µm²/Image). **(D)** Western blot analysis of total/cleaved caspase-3/8/9, TRAF2, phosho(p)/total TAK1, MKK4, JNK, NF-κB, pLKB1 and Bcl2 in IRG +/- MO3.13-A2-Luc cells upon 4 h treatment with TRAIL. Experiment was run in two separate blots each including Scr samples. Each HIT was shown in comparison to Scr sample analyzed in the same blot (Blot1: Scr, siSTK11, siKCNH8 and siABCA2; Blot2: Scr, siSLC1A3 and siCHRNA1). Representative data of (A & D) two, (B-C) three independent experiments. Values represent the mean ± SD. P-value was calculated using paired two-tailed Student’s t-test (* = p < 0.05, ** = p < 0.01, *** = p < 0.005, **** = p < 0.001).

Ligation of TRAIL-R2 with TRAIL induces apoptotic signaling through DISC (death-inducing signaling complex) formation and triggering the cleavage and activation of caspase-8.^46^ TRAIL-R2 downstream signaling can also activate pathways with dual roles on apoptosis such as JNK and NF-κB depending on the cellular context.^47^ To investigate the involvement of the IRGs on TRAIL-R2 downstream signaling, we performed Western Blot analysis on TRAIL-treated or untreated IRG proficient and deficient MO3.13-A2-Luc cells (Figure 5D). In accordance with real-time apoptosis assay (Figure 5C), we observed an increased cleavage of caspase-8 and caspase-3 after IRG silencing in MO3.13 oligodendrocytes following TRAIL treatment as an indicator for the activation of the extrinsic apoptosis pathway (Figure 5D). Only KCNH8 deficient cells showed similar caspase cleavage compared to KCNH8 proficient cells in WB analysis potentially due to early time point of analysis (2 h). This also aligns with the results of the real-time cytotoxicity assay (Figure 5C), where KCNH8 deficient cells displayed a later apoptosis induction upon TRAIL treatment compared to other IRG-deficient cells. Except for SLC1A3, the other IRGs regulated caspase-9 cleavage as demonstrated by pronounced caspase-9 activation after TRAIL exposure upon silencing of these IRGs. This indicates their role in the regulation of the intrinsic apoptosis pathway. We observed a particularly strong caspase-3/-8/-9 cleavage in STK11 deficient cells which may be related to elevated DR5 surface expression in these cells (Figure 5B).

A key driver of apoptosis signaling is c-Jun N-terminal kinase 1 (JNK1). JNK1 activation drives apoptosis through several mechanisms: transcriptional activation of pro-apoptotic genes via c-Jun/AP-1^48^, by inactivating anti-apoptotic Bcl-2 family members^49^, by inducing mitochondrial dysfunction^50^ and by enhancing caspase activation^51^. IRG-deficient MO3.13 oligodendrocytes showed elevated phosphorylated JNK1 levels in both TRAIL treated and untreated cells, except for STK11-deficient cells, where JNK1 activation was only observed upon TRAIL treatment (Figure 5D). Thus, all IRGs repress JNK1 activation during TRAIL mediated death receptor signaling. JNK is primarily phosphorylated by mitogen-activated protein-kinase kinases MKK4 or MKK7, which can be activated by multiple upstream MAP3Ks. An important, but not the only one among them is TAK1.^52, 53^ We analyzed the capacity of the five IRGs to inhibit the activation of TAK1 and MKK4. KCNH8, CHRNA1 and SLC1A3 deficient MO3.13 cells elevated pMKK4 levels after TRAIL treatment. Silencing of ABCA2 and STK11 did not affect MKK4 activation, which leaves the option that repression of JNK1 by these IRGs may be mediated through MKK7 instead of MKK4 or through MKK4/7 independent pathways. TRAIL induced TAK1 activation was regulated only by ABCA2 and CHRNA1. Thus, ABCA2 and CHRNA1 controls TRAIL-induced JNK1 activation already at the level of TAK1 activation, while KCNH8 and SLC1A3 seem to regulate MKK4 activation in MO3.13 oligodendrocytes through other upstream kinases.

Signaling through death receptors such as TRAIL receptor DR5, is balanced between those elements that induce and promote apoptosis and those that restrict and regulate it, whereby the specific components that are respectively involved can differ between individual cell types. Accordingly, TRAIL signaling can cause NF-κB activation^47^ as an apoptosis-regulating mechanism. We therefore assessed whether IRG downregulation would impact on phospho-NF-κB levels in MO3.13 oligodendrocytes. We indeed observed elevated NF-κB activation upon silencing of all five IRGs, particularly after TRAIL treatment (Figure 5D). Vice versa, NF-κB signaling can promote TRAIL-R2 expression, which we also observed upon downregulation of STK11, ABCA2 and CHRNA1 (Figure 5B).^54^ NF-κB activity regulates apoptotic cell death through multiple downstream events; a major one among them is the induction of the anti-apoptotic molecule Bcl-2. We therefore studied Bcl-2 expression in MO3.13 oligodendrocytes. Interestingly, despite NF-κB upregulation, increased silencing of the five selected IRGs did not result in increased Bcl-2 levels (Figure 5D).

Another key regulator of TRAIL mediated signaling is TNF receptor-associated factor 2 (TRAF2). TRAF2 mainly acts as a pro-survival adaptor by recruiting cellular inhibitor of apoptosis 1/2 (cIAP1/2) and linear ubiquitin chain assembly complex (LUBAC) to TRAIL-R2 which inhibits pro-caspase-8 cleavage.^55, 56^ TRAF2 can also inhibit extrinsic apoptosis by directly ubiquitinylating active caspase-8 for proteasomal degradation.^57^ We consistently observed decreased TRAF2 levels after knockdown of all five IRGs cells, with the strongest reduction upon CHRNA1 knockdown by which we also observed the strongest cleaved caspase-8 level in response to TRAIL (Figure 5D).

Finally, we explored the impact of the five IRGs on activation of STK11, which encodes for the master serine/threonine kinase^58^ Liver kinase B1 (LKB1). LKB1 is a well-established tumor suppressor kinase regulating energy homeostasis, cell-cycle and apoptosis in various tumor types such as non-small cell lung cancer (NSCLC), pancreatic cancer and breast cancer through activating AMPK and p53.^59, 60^ Beyond its tumor suppressor role, LKB1 can confer metabolic and apoptotic resistance under conditions of cell stress by inhibiting JNK-dependent stress signaling in some cell types).^61^ We therefore included phospho-LKB1 in the Western Blot analysis. As expected, silencing of STK11 in MO3.13 oligodendrocytes resulted in loss of pLKB1, but neither TRAIL treatment nor silencing of any of the other four IRGs influenced pLKB1 levels in MO3.13 oligodendrocytes (Figure 5D).

Taken together, our exploratory analyses revealed that the IRGs controlled both extrinsic and intrinsic apoptotic pathways with the exception of SLC1A3, which regulated the extrinsic pathway only. All of them controlled proapoptotic signaling through repression of JNK1 activation via MKK4 dependent and independent mechanisms and through promotion of TRAF2 expression. STK11, ABCA2 and CHRNA1 further repressed TRAIL receptor expression while KCNH8 promoted an immune suppressive phenotype that inhibited the secretion of proinflammatory cytokines by cytotoxic T cells.

## Discussion

In this study, we describe genes that mediate resistance against antigen-specific cytotoxic T cells in human oligodendrocytes. Our attempt is important, since destruction of oligodendrocytes by T cells is a major cause of disease progression in multiple sclerosis.^62^ Moreover, therapies with satisfactory efficacy for this disease are still lacking^63^ and yet, no systematic attempts have been made to characterize the “landscape” of genes mediating oligodendrocyte resistance against T cells. The immune resistance genes were identified among a panel of altogether 4155 pre-selected genes. Thus, our study does not cover all potential IRGs within the human genome. The study is further limited by i) the reductionistic experimental approach of a pure T cell: oligodendrocyte cell co-culture *in vitro*, neglecting physiological conditions *in vivo*, which may be imposed by stroma cells and extracellular matrix, ii) by the use of an oligodendrocyte cell line which cannot fully represent cell biological features of primary oligodendrocytes *in situ*, and iii) by the inevitably arbitrary determination of positive and negative thresholds for defining immune resistance function. These restrictions were enforced by the technical requirements of high-throughput screening, which requires a reproducible high number of target cells with homogeneous siRNA-based gene silencing on the one hand, and by the limited numbers of human antigen-specific cytotoxic T cells that can be expanded at a time, on the other. These limitations are likely to result in the neglect of relevant immune resistance genes and in the detection of genes with potentially low clinical relevance. On the other hand, the strength of the functional screen is that it provides excellent validation of the immunoregulatory function of the identified genes *per se.* To mitigate the weakness of gene expression differences between the cultured MO3.13 oligodendrocyte line and human oligodendrocyte subsets *in situ*, and to place them in the context of multiple sclerosis, we studied the gene expression patterns of the identified IRGs in oligodendrocyte subsets in brain tissue from MS patients and controls^12^.

The primary screen revealed 133 IRGs with an immune protective effect stronger than that of the positive control *SIK3*. A large proportion of these IRGs and related transcription factors were differentially expressed in distinct brain cell subpopulations. Subsets of OLGs and ImOLGs showed differential upregulation of multiple IRGs including *KCNH8*, *ABCA2*, *SIK3*, *SLC1A3*, *COMT*, *HCN2* and *CADM4* which represent marker genes of an OLG-characteristic IRG pattern. Some of them were differentially expressed in OLG subsets of MS patients: *SLC1A3*, *ABCA2*, *CADM4* and *HCN2* were differentially upregulated, while *KCNH8*, *COMT* and *SIK3* showed reduced expression. Since ABCA2 has an established role in myelination within the healthy CNS^64^ the observed upregulation in oligodendrocytes from MS patients suggests that it may support remyelination by enhancing the cell-intrinsic resistance to autoreactive T-cell–mediated immune attack. These findings indicate a complex, cell type specific expression landscape of multiple IRGs in different subsets of brain cells which are driven by differential expression patterns of related TFs in these populations and demonstrate their differential expression in OLG subsets under conditions of MS. Numerous such transcription factors also drive oligodendrocyte myelination and maturation programs (OLIG1, OLIG2, MYRF, NKX6-2, SOX2, SOX8, SOX10, RXRA, TCF7L2, SREBF1, SREBF2). It is thus conceivable that myelination and immune resistance may be partially driven by the same gene regulatory circuits in human oligodendrocytes. Since current therapies for MS cannot exert clinically relevant effects on the inflammatory and demyelinating pathological aspects simultaneously, this transcriptional axis of combined immune resistance and remyelination might represent an avenue for future therapies. It remains speculative, whether the individual capacity for differential upregulation of OLG-associated IRGs such as *ABCA2* impacts on the onset or severity of MS, or whether *vice versa* a constitutively low expression of such IRGs in certain individuals may increase their risk to develop the disease. Off-note, limitations of snRNA-seq analysis from the post-mortem tissues^42^ must be considered when evaluating gene expression profiles. Transcripts that are localized in the cytoplasm, with low-abundance and/or low-stability are underrepresented,^42^ which can be the reason why we could only detect a very low expression or a low percentage of positive cells for the majority of IRGs such as *STK11* and *CHRNA1*.

Although the screen contained 50 annotated immune checkpoint molecules, only five of the 133 IRGs belonged to this category: *HLA-G*^24^, *SIGLEC1*^25^, *OLR1*^26^, *NT5E*^23^ and *CEACAM6*^22^. We selected additional 52 candidates for validation in secondary screening and confirmed a robust immune protective function of 32 among them. It is surprising that canonical immune checkpoint molecules overall did not achieve particularly high ranks in our screens, although (some) tumor patients who are treated with immune checkpoint inhibitors (ICIs) are at risk to develop neurological immune-related adverse events (irAEs) such as CNS demyelination^65^ and ICIs can worsen anti-CNS autoimmune response in cancer patients with preexisting MS.^66–69^ However, these cases are relatively rare and our screens revealed many other genes with a robust, but hitherto unknown immune resistance biology. This unexpected finding may be resolved when one considers that the definition of an immune checkpoint molecule is its regulatory function on the activity of interacting immune cells. ^22, 70^ In contrast, our screen also captures the intrinsic resistance of the oligodendrocytes attacked by the T cells in addition to this feature. These cells apparently possess a broader variety of yet ill-defined protective mechanisms against fully activated, attacking cytotoxic T cells, whose protective efficacy may exceed that of classic immune checkpoint molecules. Interestingly, almost all (27) of the 32 novel IRGs have membrane localizations, regulating molecular transport between the cytoplasm and the extracellular space or intracellular organelles. Our exploratory studies on the mechanistic background of cell intrinsic immune resistance landscape exemplify for four of them, *KCNH8, ABCA2, SLC1A3* and *CHRNA1,* how membrane-associated IRGs confer resistance to T-cell and TRAIL-induced apoptosis in human oligodendrocytes by driving multifaceted mechanisms and in case of KCNH8 can also regulate T cell function. A sketch of the immune protective mechanisms of all five selected IRGs is shown in figure 6.

**Figure 6:**
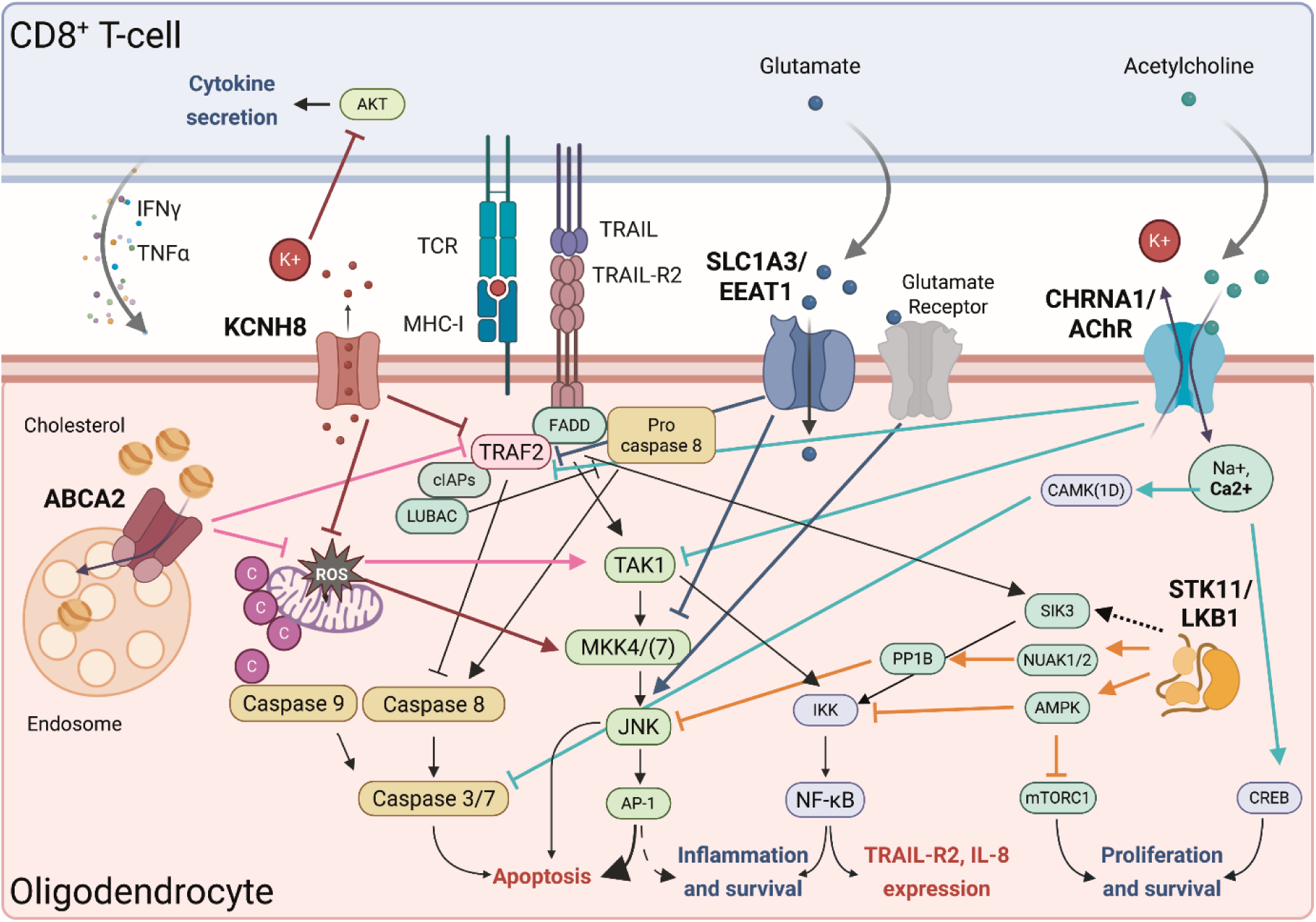
Novel immune resistance genes with multifaceted mechanisms protect oligodendrocytes from antigen-specific T-cell cytotoxicity. Summary of molecular pathways involved in the downstream signaling of STK11/LKB1, KCNH8, ABCA2, SLC1A3/EAAT1 and/or CHRNA1/AChR-mediated immune resistance. Activation of caspase-3/-8/-9, MKK4, JNK1, NF-κB and expression of TRAF2, TRAIL-R2, IL-8 was experimentally depicted in this study, whereas the potential involvement of the other molecular components was deducted from the previous studies as described and referred in the results and discussion sections above. Colors of the lines depicting the regulatory networks are matched with colors of the IRG icons. Activating and inhibitory effects are indicated by arrows and blocked lines respectively. The *figure was generated using biorender.com* (Created in BioRender. Beckhove, P. (2026) https://BioRender.com/2adzain)

*KCNH8* encodes the pore-forming subunit of a voltage-gated delayed rectifier potassium channel, which is expressed mainly in the brain and drives outward-rectifying potassium efflux.^71, 72^ Increased intrinsic K^+^ levels can suppress the activity of tumor infiltrating T cells by inhibiting Akt-signaling and IFNγ production.^73, 74^ Overexpression of K^+^ channel K_v_1.3 in T cells prevented intracellular K^+^ accumulation and K^+^-mediated immune suppression.^74^ Noteworthy, K_v_1.3 expression on myelin-specific T cells were found to be elevated in MS patients where blockade of K_v_1.3 ameliorated disease and decreased inflammation in animal models of MS.^75–77^ In this study we did not measure the change in extracellular [K^+^] upon KCNH8 knockdown in oligodendrocytes, but we hypothesize that reduced K^+^ efflux from oligodendrocytes may have caused improved T cell activity and secretion of cytokines including IFNγ and TNFα which in turn can induce JAK/STAT1/IRF1- and TNFR1-mediated apoptosis respectively. ^78, 79^ We here also demonstrate a role of KCNH8 in TRAIL-resistance as KCNH8 in MO3.13 oligodendrocytes prevented the activation of the MKK4-JNK1 axis which sensitizes cells to TRAIL-induced cytotoxicity. This is in accordance with observations that intracellular [K^+^] induces mitochondrial stress and increased ROS levels which in turn activate the MKK4-JNK1 axis (Figure 6).^80, 81^

ABCA2 (ATP-binding cassette transporter 2) is a regulator of cholesterol metabolism, a critical component of the myelin sheath formation, and most highly expressed in oligodendrocytes and neurons of the central and peripheral nervous system.^82, 83^ ABCA2 is located in the endosome/lysosome membrane where it is involved in lipid trafficking during myelination and prevents intracellular cholesterol accumulation that triggers toxic ROS production.^64, 84–88^ *ABCA2* knockout mice displayed abnormal myelin sheet formation in spinal cord and brain.^64^ The impact of ABCA2 deficiency on cholesterol-induced toxicity has been shown in macrophages previously. ^85^ Here, we show that ABCA2 protects MO3.13 oligodendrocytes from antigen-specific T cell- and TRAIL-mediated killing by both regulating TRAIL-R2 expression through suppressing NF-κB activity and by preventing proapoptotic TAK1-JNK1 signaling which can be activated by cholesterol-induced ROS accumulation during remyelination. Besides, NF-κB activation and increased TRAIL-R2 expression in ABCA2-deficient MO3.13 oligodendrocytes could be triggered by elevated intracellular cholesterol or TAK1 mediated IKK activation (Figure 6).^54, 89,90^ *SLC1A3* encodes the Excitatory amino acid transporter 1 (EAAT1), a Na^+^-dependent glutamate/aspartate transporter which is highly expressed in brain by astrocytes and oligodendrocytes, where it removes excess levels of extracellular glutamate and thereby prevents NMDA receptor-induced excitotoxicity and increases in turn intracellular glutamate concentrations.^91, 92^ Decreased EAAT1/2 expression in patients with MS and other neurodegenerative diseases such as Alzheimeŕs and Parkinsońs disease results in dysregulated glutamate homeostasis^93^ while on the contrary, it is highly upregulated in glioblastoma (GBM) cells and associated with GBM aggressiveness and immune resistance.^94,95^ Here we show that SLC1A3 reduces T cell- and TRAIL-mediated activation of the MKK4-JNK pathway and induction of apoptosis which is in accordance with previous observations that intracellular glutamate deficiency can lead to mitochondrial dysfunction and ROS accumulation,.^95, 96^, whereas elevated extracellular glutamate induce toxicity through glutamate receptor-MAPK-JNK pathway (Figure 6).^97^ *SLC1A3* silencing efficiency was observed to be lower compared to other tested IRGs, however functional data robustly demonstrated its immune-protective role in MO3.13 oligodendrocytes.

*CHRNA1* encodes the acetylcholine receptor (AChR) subunit alpha which is activated by acetylcholine^98^ to trigger neurotransmission through Na^+^/Ca^2+^ influx and calcium/calmodulin-dependent signaling in the postsynaptic cell membrane of myelinating oligodendrocytes and thereby promote proliferation and survival.^99^ *CHRNA1* mutations are associated with congenital myasthenic syndrome^100^, myasthenia gravis^101^ and amyotrophic lateral sclerosis^102^, but an involvement of *CHRNA1* in the immunopathology of MS has not yet been reported. We demonstrate that in MO3.13 oligodendrocytes CHRNA1 controls TRAF2 and TRAIL-R2 expression and TAK1-MKK4-JNK1 mediated apoptosis (Figure 6). Thereby, acetylcholine-mediated Ca^2+^ signaling could contribute to immune resistance of oligodendrocytes in MS. Our previous work revealed the immune-protective role of Calcium/Calmodulin Dependent Protein Kinase 1D (CAMK1D), where it protected multiple myeloma cells from FasL induced apoptosis through caspase-3/-7 inhibition.^16^ It will be noteworthy to investigate if AChR-triggered Ca^2+^ influx can directly activate CAMK1D in human oligodendrocytes (Figure 6).

Apart from the IRGs described above, STK11 (Liver kinase B1/LKB1) is not membrane-bound but localized within the cytoplasm. It functions as a regulator of energy homeostasis and cell growth^58^ but it’s role in cell intrinsic immune resistance is not established. We show that STK11 protects MO3.13 oligodendrocytes against T-cell/TRAIL induced apoptosis by inhibiting proapoptotic JNK1 in a TAK1-MKK4 independent manner and by suppressing TRAIL-R2 and IL-8 expression. STK11 signaling is directly linked to our positive control IRG SIK3 as the latter is a substrate kinase of STK11^15^ and similarly repressed T cell/TRAIL-induced apoptosis in MO3.13 oligodendrocytes. Yet, both LKB1 and SIK3 have various downstream and upstream regulators respectively.^103, 104^ LKB1 inhibits the activation of mTORC1 and IKK by activating AMPK ^105^ and thereby controls proliferation and the expression of apoptosis sensitizing TRAIL-R2 and IL-8.^103, 106, 107^ A recent study described a new mechanism by which LKB1 can confer drug resistance to lung cancer cells.^61^ Through phosphorylation of NUAK1/2, LKB1 can activate the PP1B phosphatase which controls JNK and inhibits mitochondrial apoptosis.^61^ Although we did not assess the direct involvement of NUAK1/2-PP1B axis in TRAIL-R downstream signaling, we hypothesize that LKB1 can potentially exert its immuno-protective role through the same mechanism in MO3.13 oligodendrocytes as we observed a strong caspase-9 cleavage and TAK1-MKK4 independent JNK activation in LKB1-deficient cells upon TRAIL treatment (Figure 6). We previously reported an immuno-protective role of the LKB1 substrate kinase SIK3, which protects tumor cells from T-cell/TNFα-induced apoptosis by promoting NF-κB nuclear translocation.^15^ In pancreatic cancer cells silencing *LKB1* did not sensitized cells to TNFα^15^, but in MO3.13 oligodendrocytes either *LKB1* or *SIK3* knockdown increased T-cell/TRAIL-induced apoptosis. Yet, we did address a direct linear TRAIL-LKB1-SIK3-JNK1 axis, as both LKB1 and SIK3 have various downstream and upstream regulators respectively.^103, 104^

Taken together, we here describe a plethora of cell intrinsic immune resistance genes in brain cell populations with a cell type selective expression pattern. Our study demonstrates how differential expression of IRGs in human oligodendrocytes exploit multifaceted mechanisms involving metabolic, potassium, cholesterol, glutamate and acetylcholine homeostasis to drive MO3.13 oligodendrocyte immune resistance and thereby potentially contribute to the immunopathology of MS. Due to low cognate receptor expression, we did not address whether IRGs confer resistance to FasL or TNFα-mediated cytotoxicity. As Fas and TNFR downstream signaling share components with TRAIL-R^108^, the involvement of IRGs in FasL and TNFα-induced oligodendrocyte apoptosis still needs to be elucidated. Notably, all five IRGs described here in more detail, converged on JNK suppression to exert their immuno-protective role in MO3.13 oligodendrocytes. This is interesting as the JNK pathway plays a peculiar role in apoptosis regulation of oligodendrocytes under stress conditions.^109–111^ However, the immune-protective role of IRGs described here still needs to be evaluated in *in vivo* models of MS such as experimental autoimmune encephalomyelitis (EAE).^112^ EAE induction in oligodendrocyte-specific IRG knockout or overexpressing mice could be a useful model to elucidate their impact in MS susceptibility and severity, and to explore potential avenues for their therapeutic targeting.

## Materials and Methods

### Cell lines

Human oligodendrocytic hybrid cells MO3.13 (Tebubio, CLU301-P, source: female) were cultured in high glucose DMEM without sodium pyruvate (Merck, #D5796) supplemented with 10% FCS, 100 U/ml penicillin G and 100 μg/ml streptomycin (Merck, #P4333) at 37 °C in a humidified atmosphere under 5% CO_2_. MO3.13-A2 cells were generated after transfection with a customized HLA-A2/Geneticin (G418) resistance gene expression plasmid (Genscript) and FACS sorting. MO3.13-A2-Luc cells were generated after transduction of MO3.13-A2 cells with lentiviral particles encoding firefly luciferase and GFP (GenTarget Inc – Amsbio, #LVP020 (CMV-Luciferase (firefly)-2A-GFP (Puro)) and FACS sorting.

### Flu antigen-specific CD8^+^ T cells

Flu antigen-specific T cells (FluTC) were generated as previously described.^20^ Briefly, peripheral blood mononuclear cells (PBMCs) were isolated from buffy coats of HLA-A2^+^ healthy donors by density gradient centrifugation (Biocoll, Merck, #L6715). On day 0, total CD8^+^ T cells were sorted from PBMCs by magnetic separation (CD8^+^ T Cell Isolation Kit, Miltenyi, #130–096-495) according to the manufactureŕs instructions and expanded in the presence of HLA-A2-matched Influenza (flu) peptide (GILGFVFTL, ProImmune, #P007-0A-E) for 14 days. The autologous CD8 negative fraction was irradiated with 60 Gray (IBL 437C Blood Irradiator) and used for 1 week as feeder cells which were then substituted by irradiated T2 cells. On day 1 and day 8, 100 U/ml IL-2 (Clinigen, #PZN-16771811) and 5 ng/ml IL-15 (R&D Systems, #247-ILB/CF) were added to the expansion. The percentage of FluTC was determined by pentamer staining (GILGFVFTL-APC, ProImmune #F007-4A-E) on day 7 and day 14 via flow cytometry (BD FACSLyric™, BD Biosciences). After antigen-specific expansion (ASE), 1 × 10^6^ FluTC were sorted by FACS and expanded further for 14 days by using a modified version of the Rosenberg protocol (rapid expansion protocol (REP)).^113^ Briefly, on day 0, sorted FluTC were cultured in 150 ml expansion media (50% AIM-V (Thermo Fisher Scientific, #12055091), 50% CLM (CLM: RPMI-1640 (Thermo Fisher Scientific, #21875091), 10% human AB serum (Valley Biomedical, #HP1022HI), 1% HEPES (Merck, #H0887), 100 U/ml penicillin G, 100 μg/ml streptomycin and 0,01% β-Mercaptoethanol (Thermo Fisher Scientific, #31350–010) supplemented with 30 ng/ml anti-CD3 (Clone: OKT3, eBioscience, #14–0037-82), 3000 U/ml IL-2 and 2 × 10^8^ irradiated allogeneic PBMCs from 3 different healthy donors. On days 5, 7 and 11, IL-2 was replenished with fresh expansion media. On day 14, cells were collected, characterized by FACS and frozen in aliquots of 5 × 10^6^ in freezing media A (60% AB serum and 40% RPMI-1640) and B (80% AB serum and 20% DMSO (Merck, #D260)). On the day of usage, FluTC were thawed at least 4 h prior to the co-culture or activation and cultured in CLM at a concentration of 1 × 10^6^ cells/ml.

### Reverse small interfering RNA (siRNA) transfection

For siRNA transfections, RNAiMAX (Thermo Fisher Scientific, #13778150) was used as described previously.^15^ For the primary and the secondary screens siGENOME siRNAs (Dharmacon^TM^; Horizon Discovery) were used (Figure 1&2). To further exclude potential off-target effects of siRNAs, we used pool of 30 siRNAs (siTOOLs) to downregulate each IRG in oligodendrocytes for functional assays (Figures 4&5). Briefly, 200 µl of 250 nM (for siRNAs from Dharmacon/Horizon) or 50 nM (for 30 pool siRNAs from siTOOLs) of siRNA solution was added per well of a 6-well plate. 4 µl of RNAiMAX transfection reagent was diluted in 200 µl final volume of RPMI-1640 (Sigma-Aldrich/Merck, #R8758) and incubated for 10 min at RT. Afterwards, 400 µl of additional RPMI was added and a total of 600 µl of RNAiMAX-RPMI mix was added to the siRNA-coated well and incubated for 30 min at RT. 2 × 10^5^ MO3.13 oligodendrocytes were resuspended in 1,2 ml of antibiotic-free DMEM culture medium supplemented with 10% FCS and seeded into the siRNA-RNAiMAX containing wells and incubated for 72 h at 37 °C. For 96-well and 384-well plate transfection, the aforementioned protocol was proportionally scaled down.

### Luciferase-based cytotoxicity assay

MO3.13-A2-Luc cells were reverse transfected with the indicated siRNA. After 72 h, cells were loaded with 0,01 µg/ml flu-peptide (GILGFVFTL, Proimmune, #P007-0A-E). After 1 h of incubation, the pulsing medium was removed and either appropriate amount of FluTC or CLM was added to MO3.13-A2-Luc cells for the cytotoxicity and viability settings respectively. After 20 h of co-culture, the supernatant was removed and the remaining luciferase activity was measured using the multimode microplate reader Spark^TM^ 10M (TECAN) as described previously.^15^ Luciferase activity is proportional to the amount of remaining live tumor cells. When indicated, raw data were normalized to negative controls.

### RNAi screening

The primary RNAi screening was conducted as described,^14, 15, 16^ using a sub-library of the genome-wide siRNA library siGENOME (Dharmacon^TM^; Horizon Discovery) comprising 4155 genes (Supp. table 1). In short, each well contained a pool of four non-overlapping siRNAs (SMARTpool) targeting the same gene. Positive and negative siRNA controls were added to each white 384-well plate (Greiner Bio-One, #781073). MO3.13-A2-Luc cells were loaded with flu-peptide and co-cultured with FluTC as described in the “luciferase-based cytotoxicity assay” section. Read-out was performed using the multimode microplate reader Spark^TM^ 10M (TECAN) with an integration time of 100 ms. Raw relative luminescence units (RLU) from the primary screen were processed using the cellHTS2 package in R/Bioconductor. Values from both conditions were quantile normalized against each other using the aroma.light package in R. Differential scores (cytotoxicity vs viability) were calculated using the locally estimated scatterplot smoothing (LOESS) local regression fitting. The thresholds for HIT calling were set according to the immune resistance gene (SIK3) which served as positive control and viability controls (UBC and cell death). For the secondary screening, a customized deconvoluted siRNA library consisting of pool of 4 non-overlapping siRNAs as well as single siRNAs targeting remaining selected 52 hits from the primary screen was used to transfect MO3.13-A2-Luc cells in 96-well plates. 72 h post transfection, cells were pulsed with flu-peptide and co-cultured either with FluTC or cultured in T cell media as performed in the primary screen. RLU values from cytotoxicity and viability settings were measured after 20 h of co-culture and normalized to Scr control.

### RNA Isolation, Reverse Transcription, quantitative RT-PCR and PCR

Total RNA was extracted from cultured cells using RNeasy Mini kit (Qiagen, #74106) and 1000 ng of RNA was transcribed using the QuantiTect reverse transcription kit (Qiagen, #205313) according to the manufactureŕs protocol. For real-time quantitative PCR (RT-qPCR) 10 ng of template cDNA, 2x QuantiFast SYBR Green PCR mix (Qiagen, #204056) and 300 nM of gene-specific primer mix (MBP fw: AGG GAT TCA AGG GAG TCG AT, MBP rev: GGG TGA TCC AGA GCG ACT AT; STK11 fw: TTG TTT GGT TGG TTC CAT TTT, STK11 rev: CAG CAG CTC CCA AAC ACC; KCNH8 fw: ATG GAG AGG GAA GAC AAC AGC C, KCNH8 rev: GCG TGA AGT ACA GAG CGG CAA T; ABCA2 fw: AAG CAC CTG CAG TTT GTC AG, ABCA2 rev: GGG ACC AGG TAG TTG AGC AT; SLC1A3 fw: GGT TGC TGC AAG CAC TCA TCA C, SLC1A3 rev: CAC GCC ATT GTT CTC TTC CAG G; CHRNA1 fw: CCC CAT CTA CCA GTC CTA AAG A, CHRNA1 rev: TTT TTG ATG CAG TTT GCA TTT T; CD274 (PD-L1) fw: TGC CGA CTA CAA GCG AAT TAC TG, CD274 (PD-L1) rev: CTG CTT GTC CAG ATG ACT TCG G, Merck) were used per 20 μL reaction and each sample was prepared in triplicates. Reactions were run using QuantStudio 3 (Thermo Fisher Scientific). Expression of the target genes was normalized to the expression of the reference gene (β-actin fw: CCT CGC CTT TGC CGA TCC, β-actin rev: GCG CGG CGA TAT CAT CAT CC) and the analysis was performed using comparative Ct method (ΔΔCT). The results were shown as fold change compared to the control siRNA transfected sample. For conventional PCR, samples were set up in a 25 μL volume using 2x MyTaq HS Red Mix (Bioline, #BIO-25048), 500 nM of gene-specific primer mix and 100 ng of template cDNA. The PCR program was set as follows: 95 °C for 3 min, 35 cycles of 3 repetitive steps of denaturation (95 °C for 15 s), annealing (60 °C for 20 s) and extension (72 °C for 15 s), and a final step at 72 °C for 7 min. PCR products were run on a 2% agarose gel and visualized using a UV documentation system (ChemiDoc^TM^, Bio-Rad).

### ELISA and Luminex assay

Supernatants of FluTC and IRG-proficient/-deficient MO3.13-A2-Luc co-cultures were analyzed by multiplex cytokine assay (MILLIPLEX MAP Human Cytokine/Chemokine Magnetic Bead Panel, 29-plex, Merck, #HCYTMAG-60K-PX29) (Figure 4C, Supp. Fig 3B-C). The assay was performed according to the manufactureŕs instructions and samples were measured using the MAGPIX Luminex instrument (Merck). Supernatants of FluTC-MO3.13-A2-Luc co-cultures and anti-CD3/CD28 activated FluTC were further analyzed for the detection of IFNγ (Human IFN-γ ELISA Set, BD OptEIA, #555142), TNFα (Human TNF ELISA Set, BD OptEIA, #555212), Granzyme B (Human Granzyme B ELISA development kit, Mabtech, #3485-1H-20) and TRAIL (human TRAIL/TNFSF10 DuoSet ELISA, R&D Systems, #DY375-05) (Supp. Fig. 1B, Supp. Fig. 3D). Experiments were performed according to the manufactureŕs instructions. Absorbance was measured at λ = 450 nm, taking λ = 570 nm as reference wavelength using the Spark 10M multimode microplate reader (TECAN). Soluble Fas ligand (FasL) was detected by multiplex cytokine assay (MILLIPLEX Human CD8^+^ T Cell MAGNETIC Premixed 17 Plex Kit, Merck, #HCD8MAG15K17PMX) (Supp. Fig. 3D).

### Western blot analysis

Western Blot was performed as described previously.^114^ The harvested pellets of cells were washed once with PBS and were resuspended in 40 µl of Cell Signaling Lysis Buffer (Merck Millipore, Cat#43-040) supplemented with protease and phosphatase inhibitors (Protease Inhibitor Cocktail Set III, EDTA-Free, Merck, #539134-1ML, 1:100; Phosphatase Inhibitor Cocktail 3, Merck Millipore, Cat#P0044-1ML, 1:100). The samples were then incubated in the fridge for 15 min under continuous rotation. Samples were centrifuged for 15 min at 17,000xg at 4 °C and the protein-containing supernatants were collected as the cell lysates. The protein concentration of cell lysates was determined using the Pierce^TM^ BCA Protein Assay Kit (Thermo Fisher Scientific, #23225) following the manufactureŕs instructions. Cell lysates (40 µg protein per sample) were incubated with NuPAGE^TM^ LDS Sample buffer (Thermo Fisher Scientific, Cat#NP0007) containing 10% β-Mercaptoethanol (Roth, Cat#4227.1) for 10 min at 70 °C. The samples were then loaded on Novex^TM^ NuPAGE^TM^ 4-12% Bis-Tris Protein-Gels (Thermo Fisher Scientific, Cat#NP0321BOX, NP0336BOX) in Mini Gel Tanks (Thermo Fisher Scientific) in the presence of NuPAGE^TM^ MOPS SDS Running Buffer (Thermo Fisher Scientific, Cat#NP0001) and the proteins were separated for 15 min at 80 V, followed by approximately 90 min at 120 V. The PageRuler^TM^ Prestained Protein Ladder (Thermo Fisher Scientific, Cat#26616) was loaded to each blot. After SDS-PAGE, the separated proteins were blotted on a nitrocellulose membrane with a pore size of 0.45 μm using the Trans-Blot® Turbo^TM^ RTA Mini 0.45 µm Nitrocellulose Transfer Kit (Bio-Rad, Cat#1620115). The transfer was performed at the Trans-Blot® Turbo^TM^ Transfer System using the “High MW” program of Bio-Rad at 25 V extended to 30 min. Transfer efficiency was subsequently determined by staining the remaining proteins in the gel with Bio-Safe^TM^ Coomassie G-250 Stain (Bio-Rad, #161-0786) and the transferred proteins on the membrane with Ponceau S solution (Sigma‒Aldrich/Merck, #p7170-1l) following the manufactureŕs instructions. After the Ponceau S staining, the membrane was washed with TBS-T, and subsequently blocked with 1X animal-free blocking solution (AFBS, Cell Signaling, #15019L) for 2 h at RT. After two washes with TBS-T for 5 min, the membrane was incubated overnight at 4 °C under continuous rotation with 5 ml of 1X AFBS containing the primary antibodies. The following antibodies were purchased from Cell Signaling and used at 1:1000 dilution: anti-Caspase-3 (#14220), anti-Caspase-8 (#9746), anti-Caspase-9 (#9502), anti-TRAF2 (#4712), anti-Phospho-TAK1 (Ser412) (#9339), anti-TAK1 (#5206), anti-Phospho-SEK1/MKK4 (Ser257/Thr261) (#9156), anti-SEK1/MKK4 (#9152), anti-Phospho-SAPK/JNK (Thr183/Tyr185) (#4668), anti-SAPK/JNK (#67096), anti-Phospho-NF-κB p65 (Ser536) (#3033), anti-NF-κB (p65) (#6956), anti-Phospo-LKB1 (Ser428) (#3482), anti-Bcl-2 (#15071). For the detection of housekeeping genes, anti-beta Actin (Abcam, #ab8226; 1:10000) and anti-Sodium Potassium ATPase (Abcam, #ab76020; 1:20,000) antibodies were used. The membranes were washed three times with TBS-T for 10 min and were incubated with AFBS containing HRP-coupled anti-mouse IgG (Cell Signaling, #7076S, 1:4000) or HRP-coupled anti-rabbit IgG (Cell Signaling, #7074S, 1:4000) secondary antibody for 1 h at RT while shaking. Unbound secondary antibodies were removed in three washing steps with TBS-T for 10 min on the shaker. Next, the membrane was incubated with a mix of the two substrates at a ratio of 1:1 using Trident femto Western HRP Substrate (GeneTex, #GTX14698) for 1 min (housekeeping genes) or 5 min (remaining proteins) in the dark. The blots were imaged at the ChemiDoc^TM^ Imaging System (Bio-Rad).

### Flow cytometry

Flow cytometry was used for the detection of proteins expressed on the membrane of MO3.13 and T cells as published.^20^ MO3.13 cells were detached from plates using PBS-EDTA, centrifuged at 500 x g for 5 min and resuspended in FACS buffer (2 × 10^5^ cells/tube). Live cells were distinguished by using Zombie Aqua™ (BioLegend, #423102) or Zombie NIR™ Fixable Viability Kit (BioLegend, #423106) according to manufactureŕs instructions followed by blocking with Kiovig (human plasma-derived immunoglobulin, Baxter, #PZN-06587176) at a concentration of 100 µg/mL in FACS buffer (PBS, 2% FCS) for 15 min in the dark on ice. Cells were washed once in FACS buffer and incubated for 20 min on ice in the dark with either fluorophore-conjugated primary antibodies or isotype control: PE anti-CD119 (IFN-γ R α chain) (Clone: GIR-208, BioLegend, #308606); APC anti-CD120a (TNF-R1) (Clone: W15099A, BioLegend, #369906); PE anti-CD120b (TNF-R2) (Clone: 3G7A02, BioLegend, #358404); BV421 anti-CD95 (Fas) (Clone: DX2, BioLegend, #305624); APC anti-CD261 (DR4, TRAIL-R1) (Clone: DJR1, BioLegend, #307208); PE anti-CD262 (DR5, TRAIL-R2) (Clone: DJR2-4 (7-8), BioLegend, #307406); anti-human HLA-A2 (Clone: BB7.2, BD, #561341), anti-CD274 (B7-H1, PD-L1) (Clone: 29E.2A3, BioLegend, #329708). T cells were stained using the same protocol with the following antibodies or their respective isotype controls: AF-700 anti-CD3 (Clone: UCHT1, BioLegend, #300424); PerCP-Cy5.5 anti-CD4 (Clone: RPA-T4, BioLegend, #300530); V450 anti-CD8 (Clone: RPA-T8, BD Bioscience, #560347); APC-Cy7 anti-CD45RO (Clone: UCHL1, BioLegend, #304227); BV605 anti-CD62L (Clone: DREG-56, BioLegend, #304833); APC anti-PD-1 (Clone: EH12.2H7, BioLegend, #329908); FITC anti-LAG3 (Clone: 11C3C65, BioLegend, #369307); PE-Cy7 anti-TIM3 (Clone: F38-2E2, eBioscience-Thermo Fisher Scientific, #25-3109-41). Stained cells were washed once, acquired with the BD FACSLyric™ flow cytometer (BD Bioscience) and data were analyzed using FlowJo software (Tree Star).

### Generation of supernatants of polyclonally activated T cells

Non-tissue culture 6-well plates were coated with 4 µg/ml anti-CD3 antibody (Clone: OKT3, eBioscience, #14-0037-82) overnight at 4 °C. On the next day, wells were washed two times with 1 ml PBS and FluTC were seeded in CLM supplemented with 1 µg/ml of human anti-CD28 antibody (Clone: CD28.2, BioLegend, #302902) at a concentration of 1 × 10^6^ cells/ml. After 24 h activation, cell suspension was centrifuged at 1400 rpm for 10 min and activated supernatant was collected into a fresh tube for further use.

### Real-time live-cell imaging assay

Apoptosis of MO3.13-A2 was determined by real-time live-cell imaging using Incucyte® SX5 live cell imager (Sartorius). Transfected MO3.13-A2 cells were either treated with supernatant of activated FluTC or 50 ng/ml TRAIL (PeproTech, #310-04). Afterwards, Incucyte® Cytotox Red Dye was added to cells (1:4000, Sartorius, #4632). Cells were imaged for the indicated time points at a 10x magnification. Apoptosis and live cell % confluency were quantified with the Incucyte® 2020B software.

### Re-analysis of snRNA-seq dataset

Processed and annotated snRNA-seq data^12^ from the white matter of post-mortem tissue of four individuals with progressive MS and five human controls without neurological disease profiled by 10x Genomics pipeline^115^ were retrieved from the Gene Expression Omnibus (GEO) under accession number <u>GSE118257</u>. Following the authorś quality control and data processing, 17.799 nuclei and 21.581 genes were retained and resulted in 23 clusters of distinct cell types and subpopulations. These were further used for our downstream differential gene expression analysis using built-in functions in “Seurat” R package. Each cell was matched and annotated to belong to one of the 23 clusters and the data was log normalized. Significantly differentially expressed marker genes (DEGs) between different cell types or conditions were called using Seurat’s FindAllMarkers or FindMarkers function with the Wilcoxon rank-sum test. Obtained lists were intersected with either our list from the primary screen or validated in the secondary list only after passing the restricted criteria (adjusted P < 0.05, min.pct ≥ 0.25 and log2 fold change ≥ 0.25). Similarly, DEG lists were intersected with top TFs retrieved from ChEA3 to co-regulate our validated IRGs.

### Statistical analysis

For statistical analyses, GraphPad Prism software v9.0 (GraphPad Software, La Jolla, CA, USA) was used. Results are reported as mean ± SD (Standard Deviation) as indicated in the figure legends. If not stated otherwise, statistical differences between the control and the test groups were determined by using two-tailed unpaired Student’s t-test. All Incucyte-based real-time cytotoxicity assay data (Fig. 4E; 5C) were analyzed using two-tailed paired Student’s t-test. In all statistical tests, a P-value ≤ 0.05 was considered significant with * = P ≤ 0.05, ** = P ≤ 0.01, *** = P ≤ 0.001 and **** = P ≤ 0.0001. All experiments with representative images have been repeated at least twice and representative images are shown. For the cumulative data, results from at least three independent experiments were combined.

## Data availability

All data associated with this study are presented in the paper or in the Supplementary Information. Publicly available datasets are listed in the methods.

## Acknowledgements

We thank LIT flow cytometry core facility for support in T cell sorting. We thank Heiko Smetak, Irina Fink, Ursula Spirk and Marcell Kupczyk for technical support. ANM received funding from Deutsche Forschungsgemeinschaft (DFG, German Research Foundation), Walter-Benjamin Fellowship (ME 6156/1-1). AH received funding from the European Union’s Horizon 2020 research and innovation programme under the Marie Sklodowska-Curie grant agreement No 861190. MX is supported by the “SATURN3 - Spatial And Temporal Resolution of Intratumoral Heterogeneity in 3 hard-to-treat CaNcers” in the BMBF funding line joint research on “Tumor Heterogeneity, Clonal Tumor Evolution and Therapy Resistance” (01KD2206A).

## Author Contributions

ANM, AH and PB designed this study and drafted the manuscript. ANM, AH, JS, VV, JM, NS, BO, AE, HL, CYC, FS, AW, LB, DT and TM performed experiments and analyzed the data. ANM, AS, VV, NK, TM and PB designed, conducted and analyzed the HTP screen. AH conducted bioinformatic analyses. RL provided statistical guidance. SS, MX, KG, TM provided critical inputs. All the authors reviewed and approved the final manuscript.

Corresponding authors Correspondence to Philipp Beckhove.

## Supplementary information

**Supplementary figure 1:**
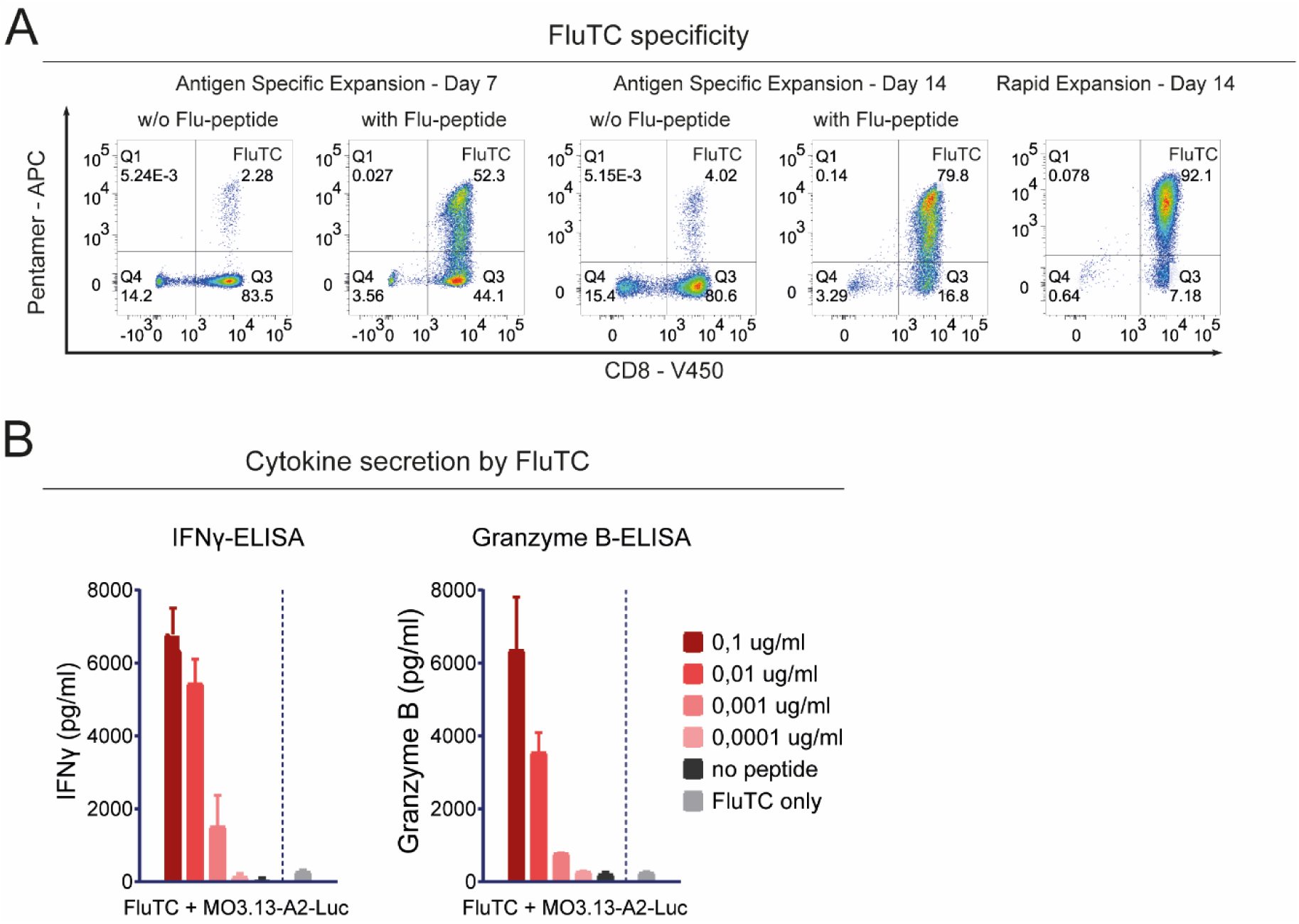
Establishment of an *in vitro* co-culture model for MS for the HTP screen, related to Fig. 1. **(A)** FACS analysis of antigen specificity of FluTC generated from HLA-A2^+^ healthy donors by repetitive expansions. All samples were gated on lymphocytes, single cells, and live cells. CD8 and Flu-Pentamer stainings were performed on days 7 and 14 of antigen-specific expansion (ASE) and day 14 of rapid expansion protocol (REP). During ASE, some CD8^+^ T cells were expanded in the presence of unpulsed feeder cells (w/o flu-peptide) as a negative control. **(B)** ELISA to determine cytokine secretion by FluTC. MO3.13-A2-Luc cells were pulsed with diluting concentrations of flu peptide and co-cultured with FluTC. After 20 h of co-culture, the supernatant of co-culture was analyzed by ELISA to detect IFNγ (left) and Granzyme B (right) secretion by FluTC.

**Supplementary figure 2:**
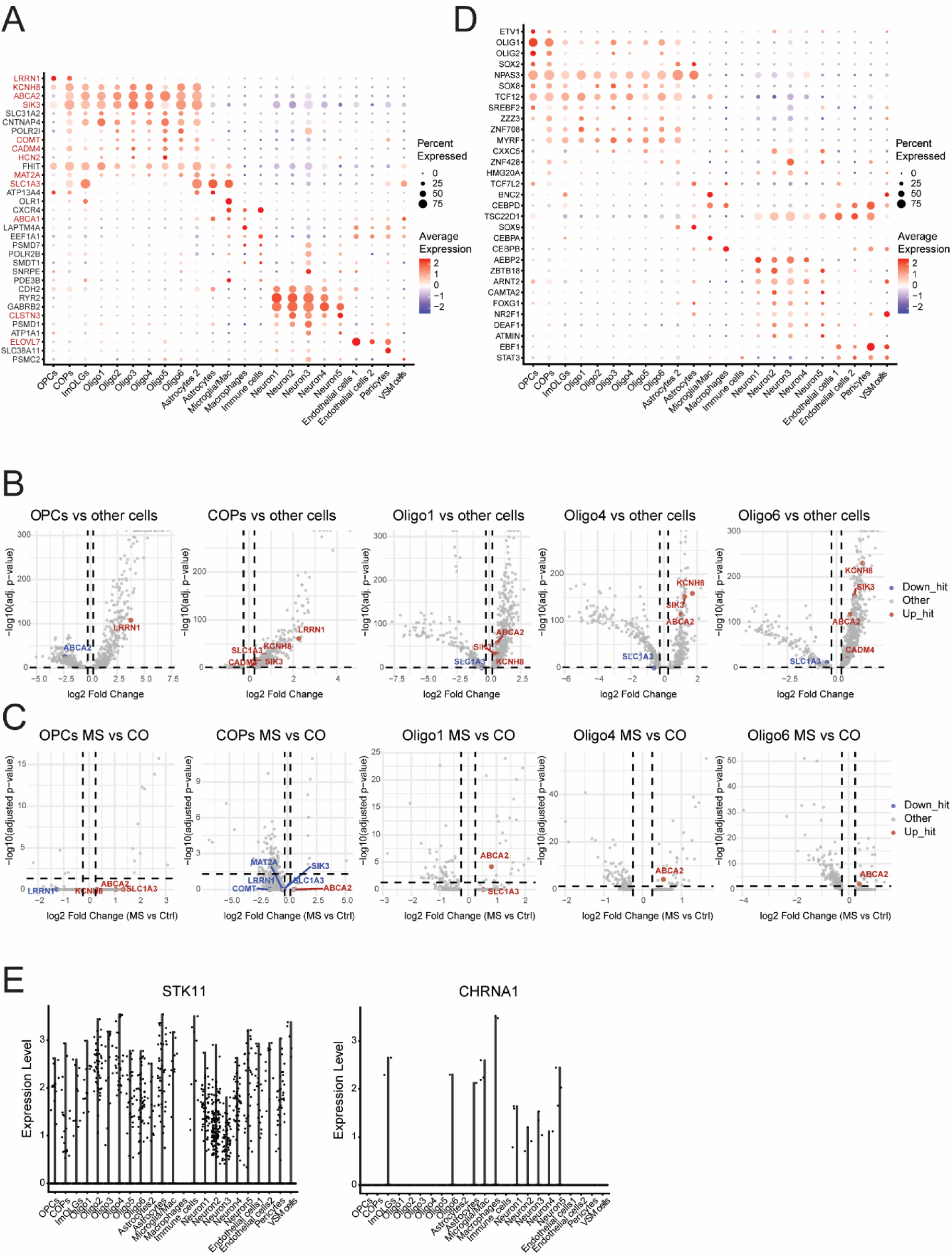
Human oligodendrocytes co-express distinct sets of IRGs with a subpopulation and MS-associated signature (related to Figure 3). Gene expression of HITs identified in the primary screen were explored among all identified populations of cells from snRNA-seq data from Jäkel *et al.* as annotated by the authors. ^12^ **(A)** Only the HITs that are among the differentially expressed marker genes of different cell clusters are shown in a dot plot where color intensity indicates average expression levels, ranging from blue (downregulated) to red (upregulated), while dot size represents the percentage of cells expressing the respective gene within every population. Validated IRGs with strong immuno-protective phenotype are marked in red. **(B-C)** Volcano plots depicting (B) DEGs of each individual oligodendrocyte subcluster versus all other cell types in MS and control samples combined and (C) DEGs of each individual oligodendrocyte subcluster from MS versus control samples. Only the IRGs with a strong immune resistance phenotype and SIK3 were highlighted. Upregulated and downregulated IRGs were marked in red and blue respectively. Horizontal dashed line indicates adjusted p-value -log10(0,05) and vertical dashed lines indicate ±log_2_(0,25). **(D)** Among the top 100 TFs predicted by ChEA3, TFs that are among the differentially expressed marker genes of different cell clusters are shown in a dot plot as in (A). **(E)** Violin plots of *STK11* and *CHRNA1* expression in all cell clusters.

**Supplementary Figure S3.**
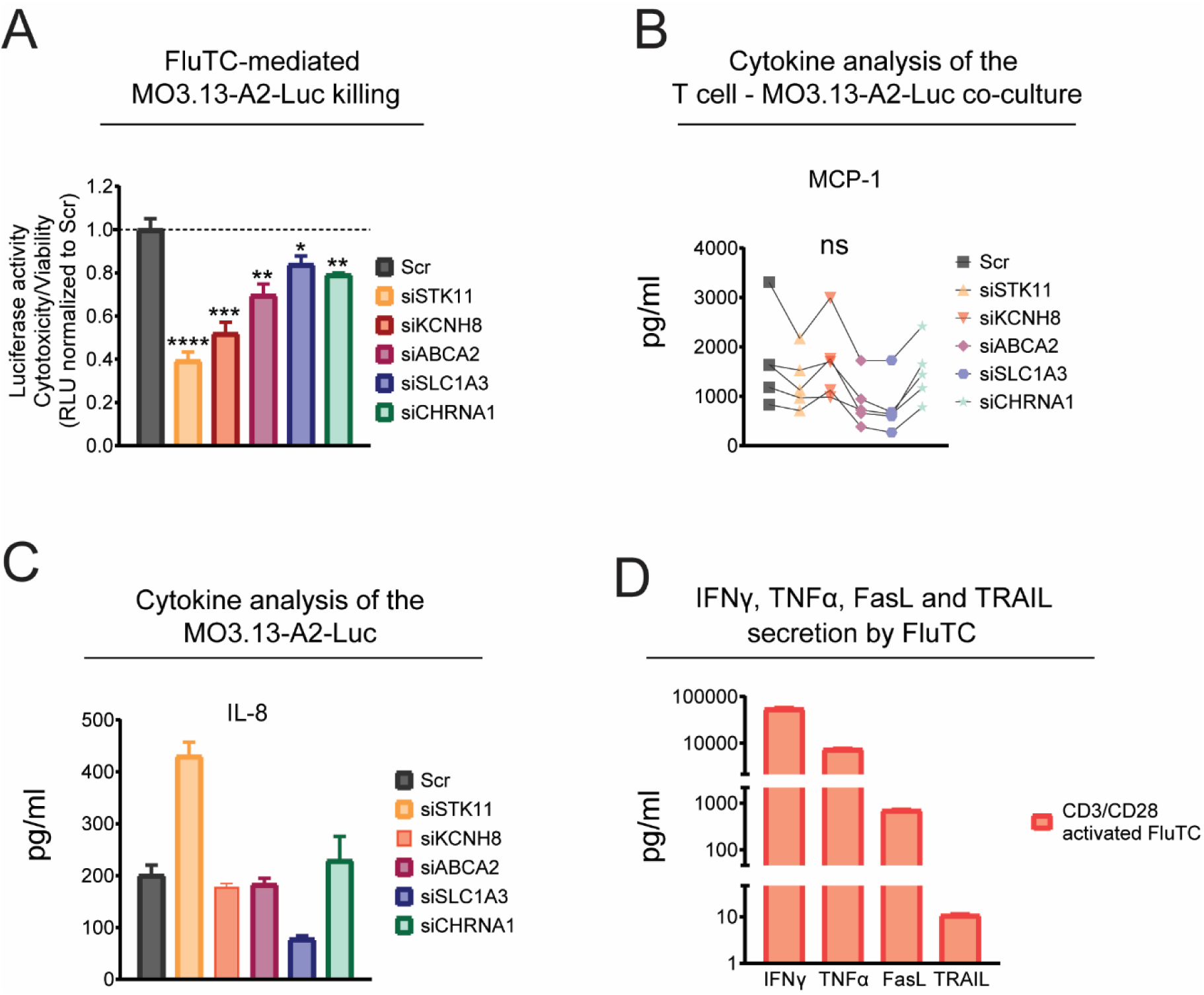
The impact of selected IRGs on T cell function and oligodendrocyte resistance against cytotoxic molecules secreted by activated T cells, related to Fig. 4. **(A)** Luciferase-based cytotoxicity assay to validate the impact of selected HITs on FluTC-mediated MO3.13-A2-Luc killing. MO3.13-A2-Luc cells were transfected with IRG-specific pool of 30 siRNA (siTOOLs) and co-cultured with FluTC as described in Fig. 1H. Graph indicates cytotoxicity/viability ratio normalized to Scr control. (B-C) Luminex-based cytokine analysis of the FluTC – MO3.13-A2-Luc cell co-cultures. MO3.13-A2-Luc cells transfected either with Scr siRNA or IRG-specific siRNA pool and (B) co-cultured with FluTC for 24 h or (C) cultured in CM. (B) MCP-1 and (C) IL-8 levels were depicted. (B) Each line represents an independent experiment; values indicate the average of (B) triplicates or (C) duplicates. (A) Representative data of at least 2 independent experiments with triplicates per sample. (A-C) Graphs show mean +/- SD. P-values were calculated using (A) two-tailed student’s t-test (B) Ratio-paired t-test. * = p < 0.05, ** = p < 0.01, *** = p < 0.005, **** = p < 0.001. **(D)** Cytokine analysis to determine IFNγ, TNFα, FasL and TRAIL secretion by anti-CD3/CD28 activated FluTC. Values represent the mean ± SD of triplicates.

**Supplementary Figure S4.**
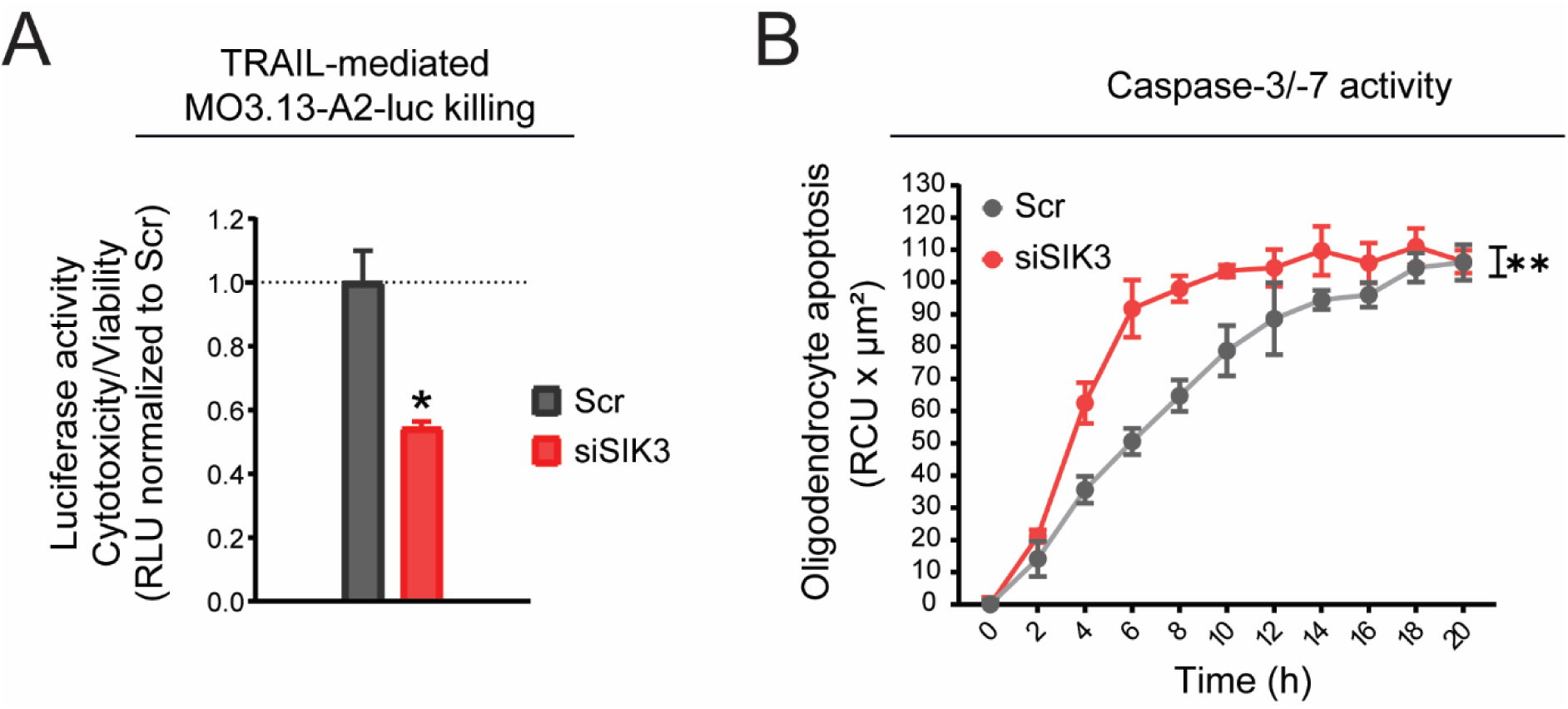
SIK3 protects MO3.13 oligodendrocytes from TRAIL-induced apoptosis, related to Fig. 5. **(A-B)** Impact of SIK3 on TRAIL-mediated MO3.13-A2-Luc killing was determined by (A) Luciferase-based cytotoxicity assay as described in supp fig. 3A and (B) real-time cytotoxicity assay as described in figure 5C. Representative data of (A) three (B) two independent experiments. Values represent the mean ± SD. P-value was calculated using paired two-tailed Student’s t-test (* = p < 0.05, ** = p < 0.01).

Supplementary Table 1 siRNA library for the primary high-throughput screen

Supplementary Table 2 Primary screen results for the top candidate 133 immune resistance genes ranked higher than SIK3

Supplementary Table 3 Performance of the well-characterized immune checkpoint molecules in the primary screen

Supplementary Table 4 Exclusion criteria for the immune resistance genes that are not followed up in the secondary screen

Supplementary Table 5 List of 32 strong immune resistance genes

Supplementary Table 6 Marker gene list of main cell types and differentially expressed genes in MS oligodendrocyte subsets compared to control individuals

Supplementary Table 7 ChEA3 analysis to identify transcription factors predicted to regulate IRG expression

## References

1. Callegari I, Derfuss T, Galli E. Update on treatment in multiple sclerosis. Presse Med. 2021;50(2):104068.

2. Filippi M, Bar-Or A, Piehl F, Preziosa P, Solari A, Vukusic S, et al. Multiple sclerosis. Nat Rev Dis Primers. 2018;4(1):43.

3. Attfield KE, Jensen LT, Kaufmann M, Friese MA, Fugger L. The immunology of multiple sclerosis. Nat Rev Immunol. 2022;22(12):734–50.

4. Becher B, Derfuss T, Liblau R. Targeting cytokine networks in neuroinflammatory diseases. Nat Rev Drug Discov. 2024;23(11):862–79.

5. Franklin RJM, Simons M. CNS remyelination and inflammation: From basic mechanisms to therapeutic opportunities. Neuron. 2022;110(21):3549–65.

6. Lubetzki C, Zalc B, Williams A, Stadelmann C, Stankoff B. Remyelination in multiple sclerosis: from basic science to clinical translation. Lancet Neurol. 2020;19(8):678–88.

7. Baecher-Allan C, Kaskow BJ, Weiner HL. Multiple Sclerosis: Mechanisms and Immunotherapy. Neuron. 2018;97(4):742–68.

8. Stinissen P, Medaer R, Raus J. Myelin reactive T cells in the autoimmune pathogenesis of multiple sclerosis. Mult Scler. 1998;4(3):203–11.

9. Cabeza-Fernandez S, White JA, McMurran CE, Gomez-Sanchez JA, de la Fuente AG. Immune-stem cell crosstalk in the central nervous system: how oligodendrocyte progenitor cells interact with immune cells. Immunol Cell Biol. 2023;101(1):25–35.

10. Gonzalez-Alvarado MN, Aprato J, Baumeister M, Lippert M, Ekici AB, Kirchner P, et al. Oligodendrocytes regulate the adhesion molecule ICAM-1 in neuroinflammation. Glia. 2022;70(3):522–35.

11. Fernandez-Castaneda A, Chappell MS, Rosen DA, Seki SM, Beiter RM, Johanson DM, et al. The active contribution of OPCs to neuroinflammation is mediated by LRP1. Acta Neuropathol. 2020;139(2):365–82.

12. Jakel S, Agirre E, Mendanha Falcao A, van Bruggen D, Lee KW, Knuesel I, et al. Altered human oligodendrocyte heterogeneity in multiple sclerosis. Nature. 2019;566(7745):543–7.

13. Schirmer L, Velmeshev D, Holmqvist S, Kaufmann M, Werneburg S, Jung D, et al. Neuronal vulnerability and multilineage diversity in multiple sclerosis. Nature. 2019;573(7772):75–82.

14. Khandelwal N, Breinig M, Speck T, Michels T, Kreutzer C, Sorrentino A, et al. A high-throughput RNAi screen for detection of immune-checkpoint molecules that mediate tumor resistance to cytotoxic T lymphocytes. EMBO Mol Med. 2015;7(4):450–63.

15. Sorrentino A, Menevse AN, Michels T, Volpin V, Durst FC, Sax J, et al. Salt-inducible kinase 3 protects tumor cells from cytotoxic T-cell attack by promoting TNF-induced NF-kappaB activation. J Immunother Cancer. 2022;10(5).

16. Volpin V, Michels T, Sorrentino A, Menevse AN, Knoll G, Ditz M, et al. CAMK1D Triggers Immune Resistance of Human Tumor Cells Refractory to Anti-PD-L1 Treatment. Cancer Immunol Res. 2020;8(9):1163–79.

17. McLaurin J, Trudel GC, Shaw IT, Antel JP, Cashman NR. A human glial hybrid cell line differentially expressing genes subserving oligodendrocyte and astrocyte phenotype. J Neurobiol. 1995;26(2):283–93.

18. Buntinx M, Vanderlocht J, Hellings N, Vandenabeele F, Lambrichts I, Raus J, et al. Characterization of three human oligodendroglial cell lines as a model to study oligodendrocyte injury: morphology and oligodendrocyte-specific gene expression. J Neurocytol. 2003;32(1):25–38.

19. Boutitah-Benyaich I, Eixarch H, Villacieros-Alvarez J, Hervera A, Cobo-Calvo A, Montalban X, et al. Multiple sclerosis: molecular pathogenesis and therapeutic intervention. Signal Transduct Target Ther. 2025;10(1):324.

20. Menevse AN, Ammer LM, Vollmann-Zwerenz A, Kupczyk M, Lorenz J, Weidner L, et al. TSPO acts as an immune resistance gene involved in the T cell mediated immune control of glioblastoma. Acta Neuropathol Commun. 2023;11(1):75.

21. Thommen DS, Schumacher TN. T Cell Dysfunction in Cancer. Cancer Cell. 2018;33(4):547–62.

22. Pinkert J, Boehm HH, Trautwein M, Doecke WD, Wessel F, Ge Y, et al. T cell-mediated elimination of cancer cells by blocking CEACAM6-CEACAM1 interaction. Oncoimmunology. 2022;11(1):2008110.

23. Allard B, Longhi MS, Robson SC, Stagg J. The ectonucleotidases CD39 and CD73: Novel checkpoint inhibitor targets. Immunol Rev. 2017;276(1):121–44.

24. Palma MB, Tronik-Le Roux D, Amin G, Castaneda S, Mobbs AM, Scarafia MA, et al. HLA-G gene editing in tumor cell lines as a novel alternative in cancer immunotherapy. Sci Rep. 2021;11(1):22158.

25. Jing W, Guo X, Wang G, Bi Y, Han L, Zhu Q, et al. Breast cancer cells promote CD169(+) macrophage-associated immunosuppression through JAK2-mediated PD-L1 upregulation on macrophages. Int Immunopharmacol. 2020;78:106012.

26. Zhang M, Fu Q, Jiang B, Yang Z, Chen Q, Chen P, et al. OLR1 is closely related to poor prognosis and immune cell infiltration in gastric cancer. Sci Rep. 2025;15(1):41611.

27. UniProt C. UniProt: the Universal Protein Knowledgebase in 2025. Nucleic Acids Res. 2025;53(D1):D609–D17.

28. Keenan AB, Torre D, Lachmann A, Leong AK, Wojciechowicz ML, Utti V, et al. ChEA3: transcription factor enrichment analysis by orthogonal omics integration. Nucleic Acids Res. 2019;47(W1):W212–W24.

29. Bujalka H, Koenning M, Jackson S, Perreau VM, Pope B, Hay CM, et al. MYRF is a membrane-associated transcription factor that autoproteolytically cleaves to directly activate myelin genes. PLoS Biol. 2013;11(8):e1001625.

30. Zhou Q, Wang S, Anderson DJ. Identification of a novel family of oligodendrocyte lineage-specific basic helix-loop-helix transcription factors. Neuron. 2000;25(2):331–43.

31. Southwood C, He C, Garbern J, Kamholz J, Arroyo E, Gow A. CNS myelin paranodes require Nkx6-2 homeoprotein transcriptional activity for normal structure. J Neurosci. 2004;24(50):11215–25.

32. Freudenstein D, Lippert M, Popp JS, Aprato J, Wegner M, Sock E, et al. Endogenous Sox8 is a critical factor for timely remyelination and oligodendroglial cell repletion in the cuprizone model. Sci Rep. 2023;13(1):22272.

33. Zhao C, Ma D, Zawadzka M, Fancy SP, Elis-Williams L, Bouvier G, et al. Sox2 Sustains Recruitment of Oligodendrocyte Progenitor Cells following CNS Demyelination and Primes Them for Differentiation during Remyelination. J Neurosci. 2015;35(33):11482–99.

34. Morrison VE, Smith VN, Huang JK. Retinoic Acid Is Required for Oligodendrocyte Precursor Cell Production and Differentiation in the Postnatal Mouse Corpus Callosum. eNeuro. 2020;7(1).

35. Zhang S, Wang Y, Zhu X, Song L, Zhan X, Ma E, et al. The Wnt Effector TCF7l2 Promotes Oligodendroglial Differentiation by Repressing Autocrine BMP4-Mediated Signaling. J Neurosci. 2021;41(8):1650–64.

36. Pruvost M, Patzig J, Yattah C, Selcen I, Hernandez M, Park HJ, et al. The stability of the myelinating oligodendrocyte transcriptome is regulated by the nuclear lamina. Cell Rep. 2023;42(8):112848.

37. Zhou Q, Anderson DJ. The bHLH transcription factors OLIG2 and OLIG1 couple neuronal and glial subtype specification. Cell. 2002;109(1):61–73.

38. Ho WY, Chang JC, Lim K, Cazenave-Gassiot A, Nguyen AT, Foo JC, et al. TDP-43 mediates SREBF2-regulated gene expression required for oligodendrocyte myelination. J Cell Biol. 2021;220(9).

39. Finzsch M, Stolt CC, Lommes P, Wegner M. Sox9 and Sox10 influence survival and migration of oligodendrocyte precursors in the spinal cord by regulating PDGF receptor alpha expression. Development. 2008;135(4):637–46.

40. Sussman CR, Davies JE, Miller RH. Extracellular and intracellular regulation of oligodendrocyte development: roles of Sonic hedgehog and expression of E proteins. Glia. 2002;40(1):55–64.

41. Choi JJ, Svaren J, Wang D. CoTF-reg reveals cooperative transcription factors in oligodendrocyte gene regulation using single-cell multi-omics. Commun Biol. 2025;8(1):181.

42. Grindberg RV, Yee-Greenbaum JL, McConnell MJ, Novotny M, O’Shaughnessy AL, Lambert GM, et al. RNA-sequencing from single nuclei. Proc Natl Acad Sci U S A. 2013;110(49):19802–7.

43. Zeis T, Enz L, Schaeren-Wiemers N. The immunomodulatory oligodendrocyte. Brain Res. 2016;1641(Pt A):139–48.

44. Tanner DC, Campbell A, O’Banion KM, Noble M, Mayer-Proschel M. cFLIP is critical for oligodendrocyte protection from inflammation. Cell Death Differ. 2015;22(9):1489–501.

45. Lopez-Gomez C, Fernandez O, Garcia-Leon JA, Pinto-Medel MJ, Oliver-Martos B, Ortega-Pinazo J, et al. TRAIL/TRAIL receptor system and susceptibility to multiple sclerosis. PLoS One. 2011;6(7):e21766.

46. Montinaro A, Walczak H. Harnessing TRAIL-induced cell death for cancer therapy: a long walk with thrilling discoveries. Cell Death Differ. 2023;30(2):237–49.

47. Cardoso Alves L, Corazza N, Micheau O, Krebs P. The multifaceted role of TRAIL signaling in cancer and immunity. FEBS J. 2021;288(19):5530–54.

48. Becker EB, Howell J, Kodama Y, Barker PA, Bonni A. Characterization of the c-Jun N-terminal kinase-BimEL signaling pathway in neuronal apoptosis. J Neurosci. 2004;24(40):8762–70.

49. Lei K, Davis RJ. JNK phosphorylation of Bim-related members of the Bcl2 family induces Bax-dependent apoptosis. Proc Natl Acad Sci U S A. 2003;100(5):2432–7.

50. Heslop KA, Rovini A, Hunt EG, Fang D, Morris ME, Christie CF, et al. JNK activation and translocation to mitochondria mediates mitochondrial dysfunction and cell death induced by VDAC opening and sorafenib in hepatocarcinoma cells. Biochem Pharmacol. 2020;171:113728.

51. Ray RM, Jin S, Bavaria MN, Johnson LR. Regulation of JNK activity in the apoptotic response of intestinal epithelial cells. Am J Physiol Gastrointest Liver Physiol. 2011;300(5):G761–70.

52. Dhanasekaran DN, Reddy EP. JNK signaling in apoptosis. Oncogene. 2008;27(48):6245–51.

53. Zeke A, Misheva M, Remenyi A, Bogoyevitch MA. JNK Signaling: Regulation and Functions Based on Complex Protein-Protein Partnerships. Microbiol Mol Biol Rev. 2016;80(3):793–835.

54. Ravi R, Bedi GC, Engstrom LW, Zeng Q, Mookerjee B, Gelinas C, et al. Regulation of death receptor expression and TRAIL/Apo2L-induced apoptosis by NF-kappaB. Nat Cell Biol. 2001;3(4):409–16.

55. Albini A, Di Paola L, Mei G, Baci D, Fusco N, Corso G, et al. Inflammation and cancer cell survival: TRAF2 as a key player. Cell Death Dis. 2025;16(1):292.

56. Lafont E, Kantari-Mimoun C, Draber P, De Miguel D, Hartwig T, Reichert M, et al. The linear ubiquitin chain assembly complex regulates TRAIL-induced gene activation and cell death. EMBO J. 2017;36(9):1147–66.

57. Gonzalvez F, Lawrence D, Yang B, Yee S, Pitti R, Marsters S, et al. TRAF2 Sets a threshold for extrinsic apoptosis by tagging caspase-8 with a ubiquitin shutoff timer. Mol Cell. 2012;48(6):888–99.

58. Lizcano JM, Goransson O, Toth R, Deak M, Morrice NA, Boudeau J, et al. LKB1 is a master kinase that activates 13 kinases of the AMPK subfamily, including MARK/PAR-1. EMBO J. 2004;23(4):833–43.

59. Bonanno L, Zulato E, Pavan A, Attili I, Pasello G, Conte P, et al. LKB1 and Tumor Metabolism: The Interplay of Immune and Angiogenic Microenvironment in Lung Cancer. Int J Mol Sci. 2019;20(8).

60. Takeda S, Iwai A, Nakashima M, Fujikura D, Chiba S, Li HM, et al. LKB1 is crucial for TRAIL-mediated apoptosis induction in osteosarcoma. Anticancer Res. 2007;27(2):761–8.

61. Li C, Syed MU, Nimbalkar A, Shen Y, Vieira MD, Fraser C, et al. LKB1 regulates JNK-dependent stress signaling and apoptotic dependency of KRAS-mutant lung cancers. Nat Commun. 2025;16(1):4112.

62. Kirby L, Jin J, Cardona JG, Smith MD, Martin KA, Wang J, et al. Oligodendrocyte precursor cells present antigen and are cytotoxic targets in inflammatory demyelination. Nat Commun. 2019;10(1):3887.

63. Olejnik P, Roszkowska Z, Adamus S, Kasarello K. Multiple sclerosis: a narrative overview of current pharmacotherapies and emerging treatment prospects. Pharmacol Rep. 2024;76(5):926–43.

64. Mack JT, Beljanski V, Soulika AM, Townsend DM, Brown CB, Davis W, et al. “Skittish” Abca2 knockout mice display tremor, hyperactivity, and abnormal myelin ultrastructure in the central nervous system. Mol Cell Biol. 2007;27(1):44–53.

65. Oliveira MCB, de Brito MH, Simabukuro MM. Central Nervous System Demyelination Associated With Immune Checkpoint Inhibitors: Review of the Literature. Front Neurol. 2020;11:538695.

66. Coureau M, Meert AP, Berghmans T, Grigoriu B. Efficacy and Toxicity of Immune - Checkpoint Inhibitors in Patients With Preexisting Autoimmune Disorders. Front Med (Lausanne). 2020;7:137.

67. Garcia CR, Jayswal R, Adams V, Anthony LB, Villano JL. Multiple sclerosis outcomes after cancer immunotherapy. Clin Transl Oncol. 2019;21(10):1336–42.

68. Gerdes LA, Held K, Beltran E, Berking C, Prinz JC, Junker A, et al. CTLA4 as Immunological Checkpoint in the Development of Multiple Sclerosis. Ann Neurol. 2016;80(2):294–300.

69. Gettings EJ, Hackett CT, Scott TF. Severe relapse in a multiple sclerosis patient associated with ipilimumab treatment of melanoma. Mult Scler. 2015;21(5):670.

70. Juneja VR, McGuire KA, Manguso RT, LaFleur MW, Collins N, Haining WN, et al. PD-L1 on tumor cells is sufficient for immune evasion in immunogenic tumors and inhibits CD8 T cell cytotoxicity. J Exp Med. 2017;214(4):895–904.

71. Baxter DF, Kirk M, Garcia AF, Raimondi A, Holmqvist MH, Flint KK, et al. A novel membrane potential-sensitive fluorescent dye improves cell-based assays for ion channels. J Biomol Screen. 2002;7(1):79–85.

72. Zou A, Lin Z, Humble M, Creech CD, Wagoner PK, Krafte D, et al. Distribution and functional properties of human KCNH8 (Elk1) potassium channels. Am J Physiol Cell Physiol. 2003;285(6):C1356–66.

73. Vodnala SK, Eil R, Kishton RJ, Sukumar M, Yamamoto TN, Ha NH, et al. T cell stemness and dysfunction in tumors are triggered by a common mechanism. Science. 2019;363(6434).

74. Eil R, Vodnala SK, Clever D, Klebanoff CA, Sukumar M, Pan JH, et al. Ionic immune suppression within the tumour microenvironment limits T cell effector function. Nature. 2016;537(7621):539–43.

75. Wulff H, Calabresi PA, Allie R, Yun S, Pennington M, Beeton C, et al. The voltage-gated Kv1.3 K(+) channel in effector memory T cells as new target for MS. J Clin Invest. 2003;111(11):1703–13.

76. Beeton C, Wulff H, Standifer NE, Azam P, Mullen KM, Pennington MW, et al. Kv1.3 channels are a therapeutic target for T cell-mediated autoimmune diseases. Proc Natl Acad Sci U S A. 2006;103(46):17414–9.

77. Rangaraju S, Chi V, Pennington MW, Chandy KG. Kv1.3 potassium channels as a therapeutic target in multiple sclerosis. Expert Opin Ther Targets. 2009;13(8):909–24.

78. Wang Y, Ren Z, Tao D, Tilwalli S, Goswami R, Balabanov R. STAT1/IRF-1 signaling pathway mediates the injurious effect of interferon-gamma on oligodendrocyte progenitor cells. Glia. 2010;58(2):195–208.

79. Malone K, LaCasse E, Beug ST. Cell death in glioblastoma and the central nervous system. Cell Oncol (Dordr). 2025;48(2):313–49.

80. Garlid KD, Paucek P. Mitochondrial potassium transport: the K(+) cycle. Biochim Biophys Acta. 2003;1606(1-3):23–41.

81. Huo JF, Hu TX, Dong YL, Zhao JZ, Liu XJ, Li LL, et al. Synthesis and in vitro and in vivo biological evaluation of novel derivatives of flexicaulin A as antiproliferative agents. Eur J Med Chem. 2020;208:112789.

82. Davis W, Jr. The ATP-binding cassette transporter-2 (ABCA2) regulates cholesterol homeostasis and low-density lipoprotein receptor metabolism in N2a neuroblastoma cells. Biochim Biophys Acta. 2011;1811(12):1152–64.

83. Kim WS, Guillemin GJ, Glaros EN, Lim CK, Garner B. Quantitation of ATP-binding cassette subfamily-A transporter gene expression in primary human brain cells. Neuroreport. 2006;17(9):891–6.

84. Villa M, Wu J, Hansen S, Pahnke J. Emerging Role of ABC Transporters in Glia Cells in Health and Diseases of the Central Nervous System. Cells. 2024;13(9).

85. Calpe-Berdiel L, Zhao Y, de Graauw M, Ye D, van Santbrink PJ, Mommaas AM, et al. Macrophage ABCA2 deletion modulates intracellular cholesterol deposition, affects macrophage apoptosis, and decreases early atherosclerosis in LDL receptor knockout mice. Atherosclerosis. 2012;223(2):332–41.

86. Njeim R, Alkhansa S, Fornoni A. Unraveling the Crosstalk between Lipids and NADPH Oxidases in Diabetic Kidney Disease. Pharmaceutics. 2023;15(5).

87. Li K, Deng Y, Deng G, Chen P, Wang Y, Wu H, et al. High cholesterol induces apoptosis and autophagy through the ROS-activated AKT/FOXO1 pathway in tendon-derived stem cells. Stem Cell Res Ther. 2020;11(1):131.

88. Banoth B, Cassel SL. Mitochondria in innate immune signaling. Transl Res. 2018;202:52–68.

89. Damhofer H, Tatar T, Southgate B, Scarneo S, Agger K, Shlyueva D, et al. TAK1 inhibition leads to RIPK1-dependent apoptosis in immune-activated cancers. Cell Death Dis. 2024;15(4):273.

90. Li Y, Schwabe RF, DeVries-Seimon T, Yao PM, Gerbod-Giannone MC, Tall AR, et al. Free cholesterol-loaded macrophages are an abundant source of tumor necrosis factor-alpha and interleukin-6: model of NF-kappaB- and map kinase-dependent inflammation in advanced atherosclerosis. J Biol Chem. 2005;280(23):21763–72.

91. Vandenberg RJ, Ryan RM. Mechanisms of glutamate transport. Physiol Rev. 2013;93(4):1621–57.

92. Pitt D, Nagelmeier IE, Wilson HC, Raine CS. Glutamate uptake by oligodendrocytes: Implications for excitotoxicity in multiple sclerosis. Neurology. 2003;61(8):1113–20.

93. Dahlmanns M, Dahlmanns JK, Savaskan N, Steiner HH, Yakubov E. Glial Glutamate Transporter-Mediated Plasticity: System x(c)(-)/xCT/SLC7A11 and EAAT1/2 in Brain Diseases. Front Biosci (Landmark Ed). 2023;28(3):57.

94. Corbetta C, Di Ianni N, Bruzzone MG, Patane M, Pollo B, Cantini G, et al. Altered function of the glutamate-aspartate transporter GLAST, a potential therapeutic target in glioblastoma. Int J Cancer. 2019;144(10):2539–54.

95. McManus RM, Komes MP, Griep A, Santarelli F, Schwartz S, Ramon Perea J, et al. NLRP3-mediated glutaminolysis controls microglial phagocytosis to promote Alzheimer’s disease progression. Immunity. 2025;58(2):326–43 e11.

96. Weinberg F, Hamanaka R, Wheaton WW, Weinberg S, Joseph J, Lopez M, et al. Mitochondrial metabolism and ROS generation are essential for Kras-mediated tumorigenicity. Proc Natl Acad Sci U S A. 2010;107(19):8788–93.

97. Thomas GM, Lin DT, Nuriya M, Huganir RL. Rapid and bi-directional regulation of AMPA receptor phosphorylation and trafficking by JNK. EMBO J. 2008;27(2):361–72.

98. Garchon HJ. Genetics of autoimmune myasthenia gravis, a model for antibody-mediated autoimmunity in man. J Autoimmun. 2003;21(2):105–10.

99. Zhang C, Guo X, Wang Y, Zhang S, Wang Z. The role of acetylcholine and its receptors in tumor immune regulation: mechanisms and potential therapeutic targets. Mol Cancer. 2025;24(1):231.

100. Finsterer J. Congenital myasthenic syndromes. Orphanet J Rare Dis. 2019;14(1):57.

101. Chia R, Saez-Atienzar S, Murphy N, Chio A, Blauwendraat C, International Myasthenia Gravis Genomics C, et al. Identification of genetic risk loci and prioritization of genes and pathways for myasthenia gravis: a genome-wide association study. Proc Natl Acad Sci U S A. 2022;119(5).

102. Anjum F, Alsharif A, Bakhuraysah M, Shafie A, Hassan MI, Mohammad T. Discovering Novel Biomarkers and Potential Therapeutic Targets of Amyotrophic Lateral Sclerosis Through Integrated Machine Learning and Gene Expression Profiling. J Mol Neurosci. 2025;75(2):61.

103. Shackelford DB, Shaw RJ. The LKB1-AMPK pathway: metabolism and growth control in tumour suppression. Nat Rev Cancer. 2009;9(8):563–75.

104. Sun Z, Jiang Q, Li J, Guo J. The potent roles of salt-inducible kinases (SIKs) in metabolic homeostasis and tumorigenesis. Signal Transduct Target Ther. 2020;5(1):150.

105. Hezel AF, Bardeesy N. LKB1; linking cell structure and tumor suppression. Oncogene. 2008;27(55):6908–19.

106. Lee HR, Yoo SJ, Kim J, Kang SW. LKB1 Regulates Inflammation of Fibroblast-like Synoviocytes from Patients with Rheumatoid Arthritis via AMPK-Dependent SLC7A11-NOX4-ROS Signaling. Cells. 2023;12(9).

107. Miyoshi T, Yamashita K, Arai T, Yamamoto K, Mizugishi K, Uchiyama T. The role of endothelial interleukin-8/NADPH oxidase 1 axis in sepsis. Immunology. 2010;131(3):331–9.

108. Xu J. The role of tumor necrosis factor receptor superfamily in cancer: insights into oncogenesis, progression, and therapeutic strategies. NPJ Precis Oncol. 2025;9(1):275.

109. Jurewicz A, Matysiak M, Andrzejak S, Selmaj K. TRAIL-induced death of human adult oligodendrocytes is mediated by JNK pathway. Glia. 2006;53(2):158–66.

110. Wang LW, Tu YF, Huang CC, Ho CJ. JNK signaling is the shared pathway linking neuroinflammation, blood-brain barrier disruption, and oligodendroglial apoptosis in the white matter injury of the immature brain. J Neuroinflammation. 2012;9:175.

111. Kim S, Dayani L, Rosenberg PA, Li J. RIP1 kinase mediates arachidonic acid-induced oxidative death of oligodendrocyte precursors. Int J Physiol Pathophysiol Pharmacol. 2010;2(2):137–47.

112. Procaccini C, De Rosa V, Pucino V, Formisano L, Matarese G. Animal models of Multiple Sclerosis. Eur J Pharmacol. 2015;759:182–91.

113. Dudley ME, Wunderlich JR, Shelton TE, Even J, Rosenberg SA. Generation of tumor-infiltrating lymphocyte cultures for use in adoptive transfer therapy for melanoma patients. J Immunother. 2003;26(4):332–42.

114. Sanna FC, Benesova I, Pervan P, Krenz A, Wurzel A, Lohmayer R, et al. IL-2 and TCR stimulation induce expression and secretion of IL-32beta by human T cells. Front Immunol. 2024;15:1437224.

115. Zheng GX, Terry JM, Belgrader P, Ryvkin P, Bent ZW, Wilson R, et al. Massively parallel digital transcriptional profiling of single cells. Nat Commun. 2017;8:14049.

